# Gene age predicts the transcriptional landscape of sexual morphogenesis in multicellular fungi

**DOI:** 10.1101/2021.06.04.447176

**Authors:** Zsolt Merényi, Máté Virágh, Emile Gluck-Thaler, Jason C. Slot, Brigitta Kiss, Torda Varga, András Geösel, Botond Hegedüs, Balázs Bálint, László G. Nagy

## Abstract

Multicellularity has been one of the most important innovations in the history of life. The role of regulatory evolution in driving transitions to multicellularity is being increasingly recognized; however, patterns and drivers of transcriptome evolution are poorly known in many clades. We here reveal that allele-specific expression, natural antisense transcripts and developmental gene expression, but not RNA editing or a developmental hourglass act in concert to shape the transcriptome of complex multicellular fruiting bodies of fungi. We find that transcriptional patterns of genes are strongly predicted by their evolutionary age. Young genes showed more expression variation both in time and space, possibly because of weaker evolutionary constraint, calling for partially non-adaptive interpretations of evolutionary changes in the transcriptome of multicellular fungi. Gene age also correlated with function, allowing us to separate fruiting body gene expression related to simple sexual development from that potentially underlying complex morphogenesis. Our study highlighted a transcriptional complexity that provides multiple speeds for transcriptome evolution, but also that constraints associated with gene age shape transcriptomic patterns during transitions to complex multicellularity in fungi.

## Introduction

The emergence of multicellularity has been one of the most influential transitions in evolution (Knoll, 2011; Smith and Szathmary, 1995). However, while simple multicellular aggregations evolved several times and evidence is accumulating that these transitions may not have had as many genetic obstacles as originally thought (Abedin and King, 2008; Kiss et al., 2019; Nagy et al., 2018; Sebé-Pedrós et al., 2017), origins of complex multicellularity (CM) seem to be rare evolutionary events. Simple multicellularity refers to cell aggregations, colonies or filaments, whereas CM comprise 3- dimensional organisms in which not all cells are in direct contact with the environment and necessitated the evolution of mechanisms for transport, cell adhesion and complex developmental programs (Knoll, 2011). Recent research suggests that besides changes in gene content or protein sequence, the evolution of gene expression and genome regulation are also important in the transition to CM (King et al., 2003; Merényi et al., 2020; Sebé-Pedrós et al., 2018). However, much of the changes underlying to origins of CM remain unknown.

Uniquely across life on Earth, fungi show evidence for multiple evolutionary origins of CM (Nagy, 2018; Nguyen et al., 2017) as well as for transitions between simple hyphae and 3-dimensional organization during the development of sexual fruiting bodies. The latter allows real-time transcriptomic readout of changes in complexity level, which make fungi an ideal model system to investigate CM. Fungi reached highest multicellular complexity in fruiting bodies of mushroom-forming fungi (Agaricomycetes, Kües & Navarro-González, 2015; Nagy, 2018) which includes most industrially cultivated edible and medicinal mushrooms. CM fruiting bodies in the Agaricomycetes have been widely studied by transcriptomic approaches, however, the interpretation of transcriptomes has been complicated by the lack of an understanding of the general principles of transcriptome evolution. This has impeded a general synthesis on the genetics of CM in fungi. Recent studies of fruiting body development reported species-specific and conserved genes (Krizsán et al., 2019; Nguyen et al., 2017), long non-coding RNA (Kim et al., 2018a), natural antisense transcripts (Muraguchi et al., 2015; Ohm et al., 2010; Shao et al., 2017), allele specific expression (Gehrmann et al., 2018), RNA-editing (Wu et al., 2019, 2018), small RNA (Lau et al., 2018), alternative splicing (Krizsán et al., 2019), chromatin remodelling (Vonk and Ohm, 2021) as well as transcriptome conservation patterns predicted by the developmental hourglass hypothesis (Cheng et al., 2015) however, how these contribute to CM is not known.

Similarly, several genes and cellular processes have been identified in fruiting bodies. Fruiting bodies are composite structures in which structural cell types enclose reproductive ones (basidia, meiospores) into a protective environment. Basidium and spore development are evolutionarily significantly older than CM fruiting bodies. The underlying genes show up in developmental transcriptomes, which, if not properly accounted for, can blur signals for real CM-related genes. Accordingly, while some hitherto identified genes can be linked to CM functions (e.g. defense of fruiting bodies Künzler, 2018), most fruiting body-expressed genes, including those related to cell wall remodeling (Liu et al., 2021), transcriptional regulation, selective protein degradation (Krizsán et al., 2019), complex secretomes (Almási et al., 2019) may relate either to CM or more general functions.

The goal of this study was to systematically tease apart the components and driving forces of transcriptome evolution in CM fungi. To this end, we compared the transcriptomes of eight CM fungi and that of a simple yeast-like species (*Cryptococcus neoformans*). We examined natural antisense transcripts, allele-specific expression and RNA-editing and found that evolutionary age predicts, to a large extent, the behaviour of a gene in the CM transcriptome. Because the simple vegetative (hyphal) and complex reproductive (fruiting body) stages are distinct, we expect morphogenetic genes specifically expressed in the complex multicellular stage to exist (Krizsán et al., 2019), in contrast to animals and plants, where such sharp separation cannot be expected. Fruiting body transcriptomes showed a clear gene-age related stratification, allowing the separation of genes related to general sexual processes from those likely linked to sculpting CM fruiting bodies. These data will help understanding both complex multicellular and simple, sexual morphogenesis in fungi.

## Results and Discussion

### Overview of new RNA-Seq data

We present the first highly resolved developmental transcriptome data for *Pleurotus ostreatus* (oyster mushroom), one of the three most widely cultured species worldwide (Zhu et al., 2019), as well as for *Pterula gracilis*, a closely related species with a simple fruiting body morphology (Fig. 1/a). In *P. ostreatus* we sampled six developmental stages and up to four tissue types within a stage, whereas in *Pt. gracilis*, tissues could not be separated, therefore we sampled four developmental stages (Fig. S1–2). Strand-specific RNA-Seq yielded 15.9-34.0 million reads per sample (Table S1). Multidimensional scaling of the normalized transcriptome data accurately identified sample groups with biological replicates being tightly positioned together (Fig. S3). For uniformity in downstream analyses, we reanalysed data from former studies (Almási et al., 2019; Gehrmann et al., 2018; Ke et al., 2020; Krizsán et al., 2019; Liu et al., 2018; Sipos et al., 2017), yielding data for eight species in the order Agaricales, which comprises a single origin of complex fruiting body morphologies (Marisol et al., 2020; Varga et al., 2019). *P. ostreatus* and *Pt. gracilis* had 4,294 and 474 developmentally expressed genes (≥4FC), respectively, within the range reported earlier (Almási et al., 2019; Krizsán et al., 2019; Sipos et al., 2017) (Fig. S4).

**Figure 1.**
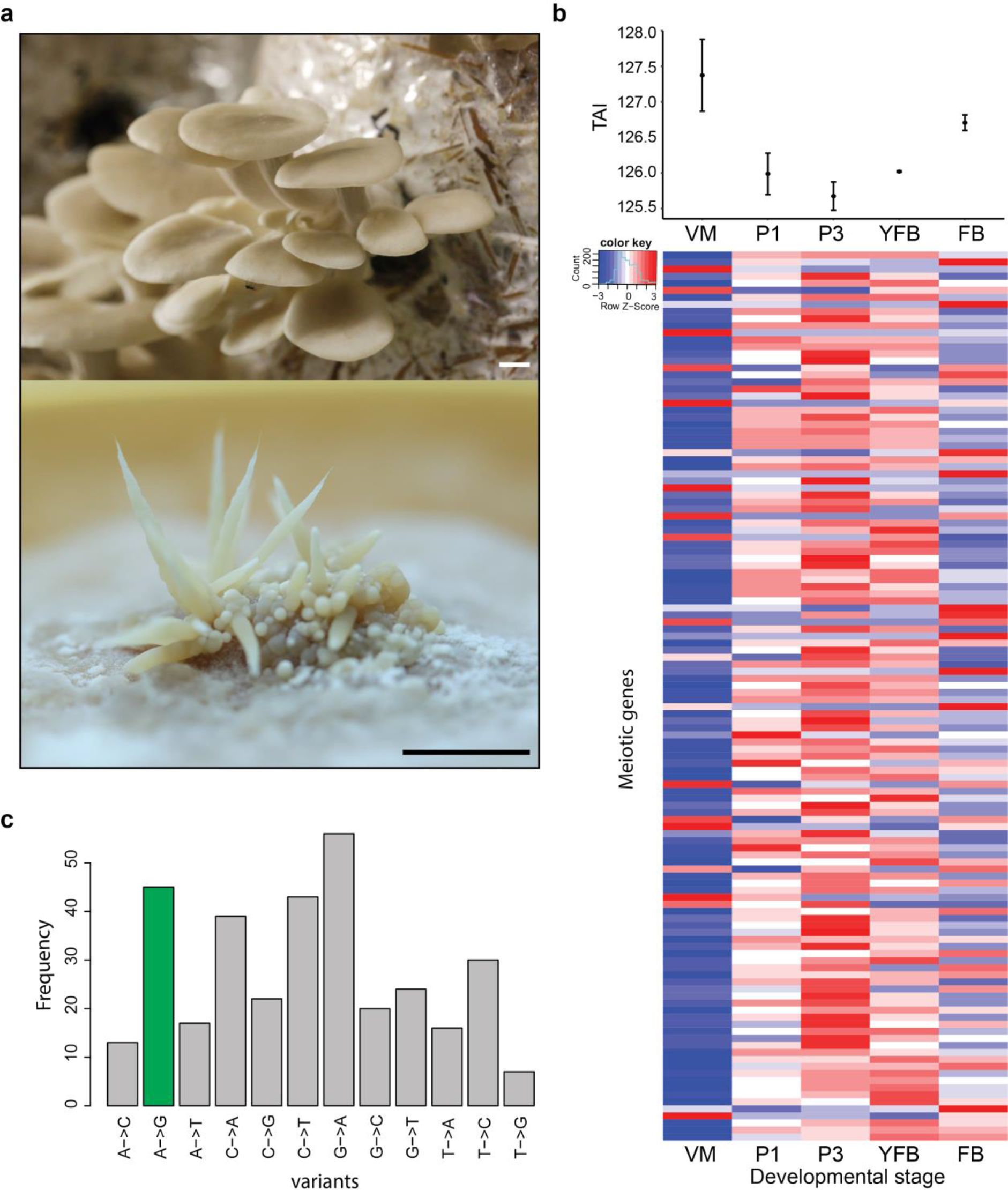
Overview of developmental transcriptomes. **a)** Fruiting bodies of *Pleurotus ostreatus* (upper) and *Pterula gracilis* (lower). **b)** Transcriptome conservation in *P. ostreatus* based on transcriptome age index (TAI) and its correlation with meiotic gene expression **c)** distribution of variant types retained in the RNA editing specific pipeline with A-to-I variants being marked with green. Abbreviations as follows: ‘VM’ vegetative mycelium; ‘P1’ stage 1 primordium; ‘P3’ stage3 primordium; ‘YFB’ young fruiting body, ‘FB’ fruiting body; ‘H’ cap (entire); ‘C’ cap trama (only the inner part, without Lamellae, or skin); ‘L’ lamellae; ‘S’ stipe; ‘V’ cuticle; ‘D’ dedifferentiated tissue of cap.

### Developmental hourglass

Previously, an hourglass concept has been proposed to explain the incorporation of genetic novelty into the developmental programs of CM eukaryotes (Domazet-Lošo and Tautz, 2010; Drost et al., 2017), including fungi (Cheng et al., 2015). This concept posits that evolutionarily older genes are expressed at mid-development, while younger genes are expressed at early and late stages (Domazet-Lošo and Tautz, 2010). To test this hypothesis, we analyzed the transcriptomes of the nine species based on transcriptome age indexes, which measure transcriptome conservation. We obtained hourglass-like patterns only in *P. ostreatus, M. kentingensis* and *A. bisporus*, whereas in other species no clear-cut signal for a developmental hourglass was found (Fig. 1/b, Fig. S5). We observed that, in each species, the highest level of transcriptome conservation coincided with mitotic/meiotic gene upregulation (Fig. 1/d), a well-known signal for cell proliferation and sporulation-related meiosis. This suggests that in fungi, transcriptome conservation patterns are merely driven by expression peaks of ultra-conserved genes related to the cell cycle, and may not reflect mechanisms for incorporating genetic novelty, as hypothesised for metazoans (Domazet-Lošo and Tautz, 2010).

### Developmental gene clusters

We found evidence for the grouping of developmentally expressed genes into hotspots in the genomes (see Methods) (Table S2, Fig. S6). Altogether 153 hotspots were detected in eight genomes; however, most of these were species-specific and not conserved across species (e.g. the luciferase cluster in the bioluminescent *A. ostoyae,* Ke et al., 2020b).

### Allele-specific expression but not RNA editing is abundant in fruiting body transcriptomes

*P. ostreatus* is a good model to distinguish the contributions of RNA editing and Allele-Specific Expression (ASE) because both parental genomes are sequenced, and sufficiently differ from each other to classify variants either as ASE (variants differing from one parental genome) or RNA editing (variants differing from both parental genomes).

Overall, 2,244,348 variants were input to the ASE analysis and were used to decide which haploid nucleus the reads originated from (Table S3). We inferred that 31.2% and 32.2% of the reads derive from PC15 and PC9 nuclei, respectively, while 36.5% of reads were not able to be assigned to either parental genomes (Table S3). This allowed us to characterize 9,948 PC15 genes (80.7% of all genes and 95.1% of expressed genes) for ASE. Similar to gene expression, allele specific expression levels showed clear stage- and tissue-specific patterns (Fig. S7–8).

At the scale of the entire genome or scaffolds, the two parental genomes expressed almost equally (Fig. S9–10), whereas at the gene level 7,793 (74,8%) of the 10,419 expressed genes were assigned as Equally Expressed genes (EE genes, hereafter) in all stages and tissue types and 2,626 genes (25.2%) was biased towards the same nucleus in all biological replicates of at least one stage or tissue (referred to as ASE genes, Fig. 2/c). Of these, 1,560 genes showed 2 < FC < 4-fold expression imbalance (hereafter referred to as S2 genes; 15%) and 1,066 (S4 genes; 10.2%) showed over four-fold difference between the two nuclei in at least one stage (averaged across replicates, Table S4). In comparison, in *A. bisporus* ASE was reported for 411 genes (~4% of the genome), perhaps due to fewer SNPs between and, accordingly, lower power in assigning reads, to parents (Gehrmann et al., 2018).

**Figure 2.**
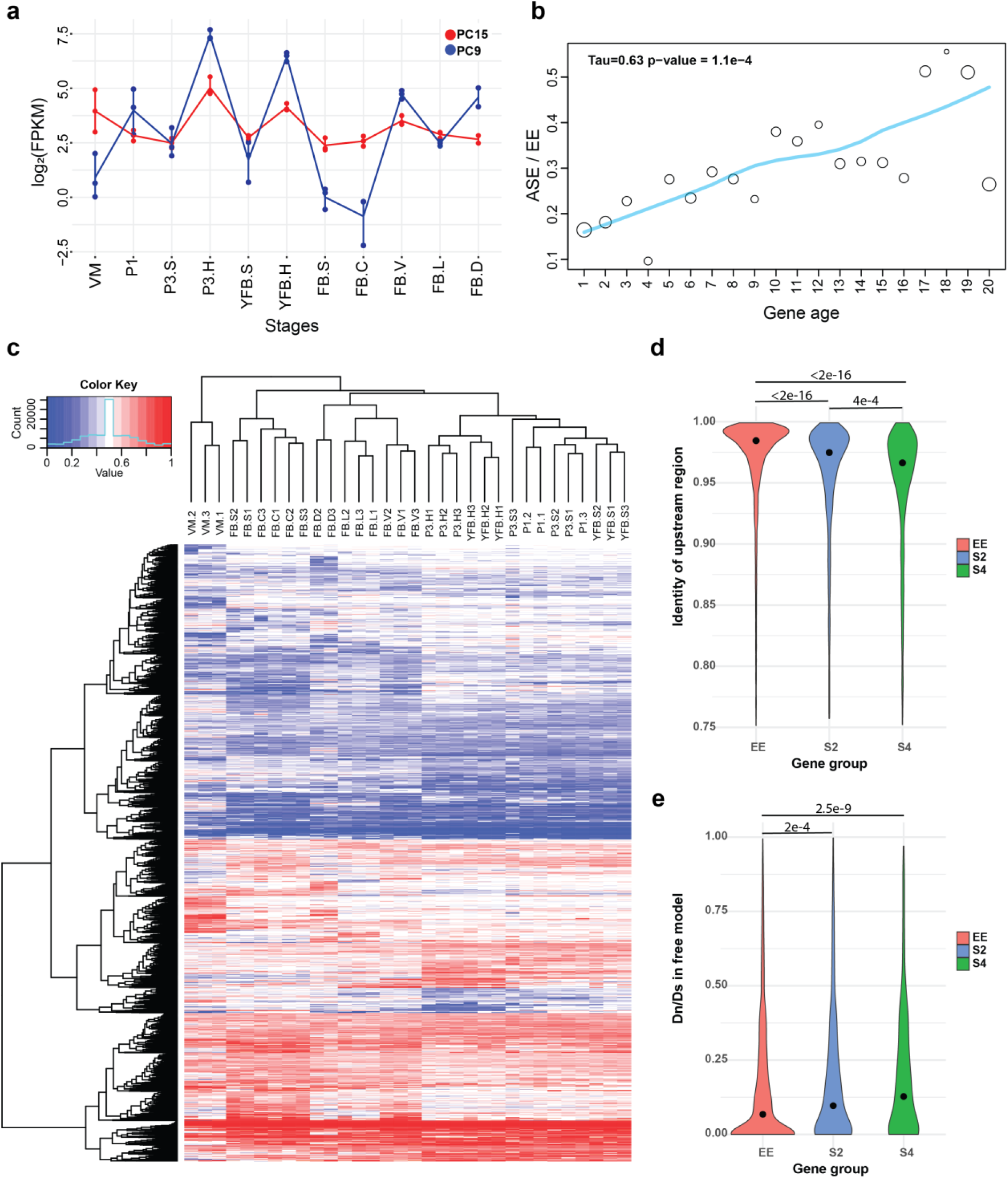
Transcriptome patterns correlate with gene age in *Pleurotus*. **a)** allele specific expression (ASE) of Mycotoxin biosynthesis protein UstYa-like domain containing protein (PleosPC15_2_159921) **b)** Proportion of ASE shows a significant increasing tendency (Mann-Kendall statistics) across gene ages. Size of circles represents the number of proteins (log10 transformed). **c)** Heatmap based on AS ratio (expression contribution of PC15 relative to the sum of PC15 and PC9) of genes that showed at least two-fold ASE in at least one stage. **d)** percent sequence identity between the upstream regions of PC15 and PC9 genes **e)** dN/dS distribution for three gene groups under the free (fb) model in CODEML. Abbreviations as follows: ‘VM’ vegetative mycelium; ‘P1’ stage 1 primordium; ‘P3’ stage3 primordium; ‘YFB’ young fruiting body, ‘FB’ fruiting body; ‘H’ cap (entire); ‘C’ cap trama; ‘L’ lamellae; ‘S’ stipe; ‘V’ cuticle; ‘D’ dedifferentiated tissue of cap.

Enrichment analysis based on InterPro domains and GO terms of ASE genes highlighted a significant overrepresentation of some well-known fruiting-related gene families (Krizsán et al., 2019), such as hydrophobins, glycoside hydrolase families, aquaporins and fungal type protein kinases (Table S5-6, Fig. S11). Notably, all of these families are volatile, characterized by frequent duplications.

In the RNA-editing pipeline (Fig. S12), 627,093 of the 1,999,221 input variants remained after filtering for extreme low frequency (< 0.1%). We chose this permissive threshold (as opposed to 1/3/10% in other protocols (Liu et al., 2017; Wu et al., 2018), to avoid discarding any signal in the early steps. As many as 546,790 sites were located in gene regions, of which 346,105 possibly corresponded to erroneous mapping around splice sites (Fig. S13), while 218,685 were retained for further analysis. Surprisingly, only 1.2% of these (2,701 sites) were consistent between at least two of the three biological replicates. After eliminating potential sequencing errors (Wilcoxon signed-rank test, p-value<0.01), we obtained 1,179 variants. Requiring at least 1% mean variant frequency in at least 1 stage left 332 potential RNA editing sites, with 6 to 62 in each variant type. (Table S7). Among these A-to-I and C-to-U transitions were not enriched (Fig. 1/c), consistent with previous Basidiomycota studies (Bian et al., 2019; Teichert, 2020). Wu et al. (2018) hypothesized that the lack of A-to-I enrichment indicates a greater diversity of editing sites and novel mechanisms of RNA editing in Basidiomycetes. To explore other explanations, we examined what, other than RNA-editing, our remaining variants could potentially correspond to. By examining motifs around the 332 sites, we detected solely an enrichment of adenines 1-2 nucleotides downstream of C-to-A sites (Fig. S14/b). However, because 69% (20 of 29) of these were within +/−20nt from the 3’ end of the last exon of genes, we think that adenine enrichment corresponds to the polyadenylation sequence. Together, we interpret these results as limited or no evidence for RNA editing, but abundant occurrence of allele-specific expression in *P. ostreatus*.

### Allele-specific expression is enriched in young genes

We next asked what mechanisms could give rise to ASE. Gehrmann et al. (2018) found that DNA methylation can explain at most 10% of ASE, which is consistent with the negligible role of gene body methylation in fungi (Montanini et al., 2014), suggesting other mechanisms. Following reports of divergent cis-regulatory alleles causing allelic gene expression imbalance (Gaur et al., 2013; Chen et al., 2016; Cowles et al., 2002; McManus et al., 2010; M. Wang et al., 2017), we hypothesised that ASE may arise from cis-regulatory divergence between nuclei of *P. ostreatus*. Indeed, upstream 1.2 kb regions, which probably comprise cis-regulatory elements, of S2 and S4 genes are significantly more different (Kruskal-Wallis with Nemenyi post hoc test *P* < 2e-16) between the two parents than those of EE genes (Fig. 2/d). This is consistent with divergent cis-regulatory elements in the same trans-regulatory cellular environment causing differential binding of transcription factors, thus causing biased gene expression in the two nuclei. Protein sequences of S4 and S2 genes are also significantly more different between the two parents (Kruskal-Wallis test with Nemenyi post hoc test *P*<2.2e-16 and 2.7e-14 Fig. S15/a) than that of EE genes, and dN/dS also showed an increased level among ASE genes (Kruskal-Wallis test with Nemenyi post hoc test *P*=2.5e-9 and 2.0e-4 Fig. 2/e, Fig. S15/b). Together, these observations indicate that ASE in *P. ostreatus* may arise from the divergence of cis-regulatory alleles, possibly in fast-evolving genes that are released from selection constraints.

A well-known group of genes under relaxed selection are evolutionarily young genes, therefore, we tested whether ASE is correlated with relative gene age in our dataset (gene ages were assigned based on time since last duplication, see Methods). ASE genes showed a strong and significant overrepresentation in the three youngest gene ages (Fisher’s exact test P=2.6e-7 - 4.8e-44), with clear trends (Fig. 2/b, Mann-Kendall test P= 1.1e-4), which together contain 51,2% of ASE genes. At the same time, they are significantly underrepresented in the three oldest age categories (gene age 1-3: *P*=2.9e-3 to 1.5e-49, Fisher’s exact test, Fig. 2/b). These observations are consistent with recently duplicated genes tolerating expression imbalance better than conserved genes, probably due to relaxed constraint (Dong et al., 2011; Gu et al., 2005; Kondrashov et al., 2002).

If genes under weak selection can tolerate expression variation, and developmental expression is considered an adaptively or neutrally arising expression variation then ASE genes and developmentally expressed genes should overlap to some extent. Indeed, half of the ASE genes (S4: 52.7% and S2: 49.1%) were also developmentally regulated (FC>4), significantly more than in EE genes (31.8%, Fisher’s Exact Test p=8.2e-58). We observed that as we move towards younger genes, the proportion of developmentally expressed ASE genes increase, as compared to non-ASE genes (Fig. S16–S17). The strongest overrepresentation of ASE genes can be observed among developmentally expressed genes that arose in the last common ancestor of *Pleurotus* (gene age 18, P_S4/EE_=7.8e-16, Fisher’s Exact Test).

### Natural Antisense Transcripts show fast turnover

Natural antisense transcripts (NATs) are abundantly transcribed from fungal genomes and can include important regulatory RNAs that influence, among others, sexual development (Donaldson et al., 2017; Donaldson and Saville, 2012; Faghihi and Wahlestedt, 2009; Kim et al., 2018a). We analysed NATs in *P. ostreatus* and *Pt. gracilis* based on strand-specific RNA-seq data, and we identified 2,043 and 763 *de novo* transcripts as NATs (Fig. S18), corresponding to 17.6% and 6.3% of protein coding genes, respectively. Lengths, exon structures and coding potentials of the assembled NATs were similar to those in earlier reports (Borgognone et al., 2019; Kim et al., 2018a; Wang et al., 2019) (Fig. S18–S19).

NATs showed developmentally dynamic expressions with NAT expression patterns reflecting stage and tissue identity, with tight grouping of biological replicates (Fig. S20). As many as 1173 (57.4%) and 126 (16.5%) NATs of *P. ostreatus* and *Pt. gracilis* were developmentally expressed, respectively. These may expand the space of developmentally regulated transcripts in fruiting bodies, thus, irrespective of their exact mechanism of action, can contribute to CM. In *P. ostreatus* we identified 166 NATs (8.1%) that showed at least 2-fold higher expression in all fruiting body stages than in vegetative mycelium, comparatively more than genes (4.8%). Such transcripts may regulate the transition from simple multicellularity in VM to complex multicellularity in FB, one of the most significant transcriptomic reprogramming events in the fungal life cycle (Krizsán et al., 2019). Kim et al. (2018) found a similar proportion (21.3%, 547 of 2,574) of lncRNA that might have a role in sexual development of *Fusarium graminearum*. Nevertheless, only 39.1% of *P. ostreatus* and 4.1% of *Pt. gracilis* of the NAT-possessing genes show developmental regulation.

The detected NATs displayed low sequence conservation. Of the 2,043 NATs of *P. ostreatus*, 1,815 (89%) showed homology in the closest sequenced species (*P. eryngii*) and only 177 (8.7%) in other species (Fig. 3). In *Pt. gracilis* only 15 (2.0 %) NATs showed homology in other species (Fig. S21). We find that low overlap with gene exons can explain the lack of sequence conservation in NATs. In *P. ostreatus*, only 596 of the 2,043 NATs (29%) showed at least 75% total exon-exon overlap. This suggests that, even if located in conserved genes, NATs mostly overlap with introns or intergenic regions, which may allow rapid sequence turnover, as reported in vertebrates (Kapusta and Feschotte, 2014). We also failed to detect conservation of sense gene identity: of the 6,232 co-orthologous genes between *P. ostreatus* and *Pt. gracilis*, only 70 showed evidence for NAT in both species. All of these observations imply low conservation of NATs at the sequence level, consistent with the view that NAT homology is detectable only among closely related species (Donaldson and Saville, 2013; Rhind et al., 2011).

**Figure 3.**
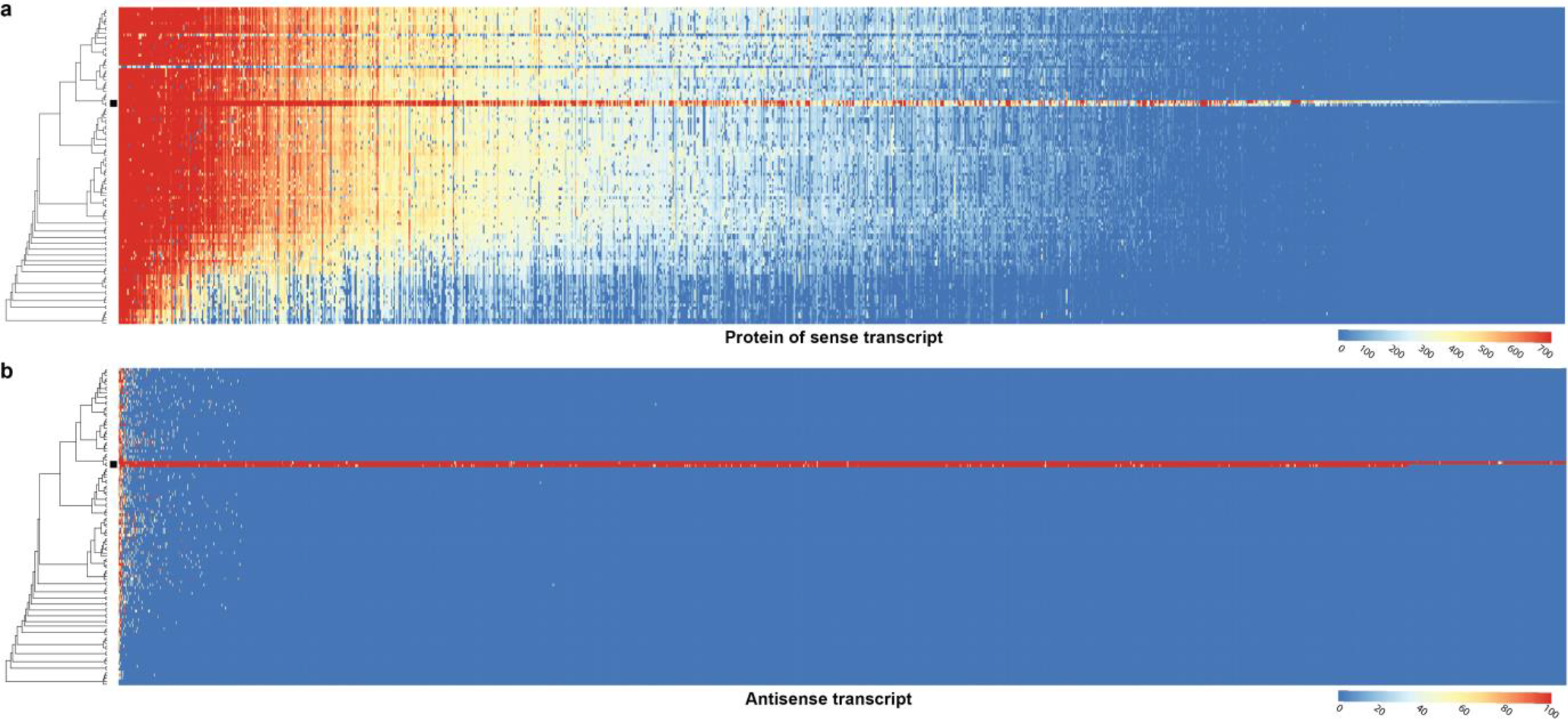
Conservation of sense genes and their antisense transcripts (NAT) in *Pleurotus ostreatus*. **a**) similarity of sense transcripts measured with −log10(e-value) from MMSeqs search **b**) mapping score of antisense transcripts based on Minimap2. Warmer colour represents a higher similarity according to the scales. Black square denotes *P. ostreatus*. Rows represents the species while columns represent the antisense query transcripts (b) or proteins from sense transcripts (a). For a larger species tree and the same plot in *Pt. gracilis*, see Figs. S29 and S21, respectively.

In *P. ostreatus* 263 NATs (12.8%) showed significant positive (Pearson r > 0.7, P < 0.05, Fig. S22 twice as many as expected by chance P<0.01, Fig. S23) while 33 showed significant negative expression correlation (Pearson r < −0.7, P <0.05, Fig. S24) with their sense genes. An enrichment of positive over negative correlation between lncRNA/gene pairs was noted previously in *Fusarium* and in *Ganoderma lucidum* (Kim et al., 2018b; Shao et al., 2017). Positively correlating pairs may be co-regulated via chromatin accessibility, or be stabilized via dsRNA formation (Donaldson and Saville, 2013), whereas negative correlation can be explained by transcriptional interference, antisense mediated chromatin remodelling or RNA masking (reviewed in Donaldson & Saville, 2012).

Together, the developmentally relevant expression, the lack of functional clues (Fig. S25) and the low conservation of NATs suggests that antisense transcription is a fast-evolving component of CM transcriptomes with potential functions in modulating gene expression. Nevertheless, as above, non-adaptive explanations should not be ruled out, such as some NATs being transcriptional noise or leakage (Dahary et al., 2005; Lloréns-Rico et al., 2016).

### Sexual and CM development recruit genes of different ages

After we asserted/contended that transcriptome patterns may be affected by gene age and constraint (or lack thereof), we examined age distributions of developmentally expressed genes. We found that developmental transcriptomes are dominated by old and young genes in all species, creating ‘U’ shaped distributions (Fig. 4/c, Fig. S26). While this shape mirrors the genome-wide gene age figures, most species displayed an enrichment of developmental expression (at fold change>4) among young genes (Fisher’s exact test, FDR corrected *P*<0.05, Fig. 4/c, Fig. S26), indicating that, like in the case of ASE, young genes have a disproportionately high share in fruiting bodies. To eliminate young genes and focus on key CM functions, we first looked for 1-to-1 orthologous genes, hereafter called ‘orthogroups’ (1 gene per species), which show developmentally dynamic expression in most species (FC>2/4, see Methods). This yielded 1,781 orthogroups, considered hereafter as conserved developmental orthogroups.

**Figure 4.**
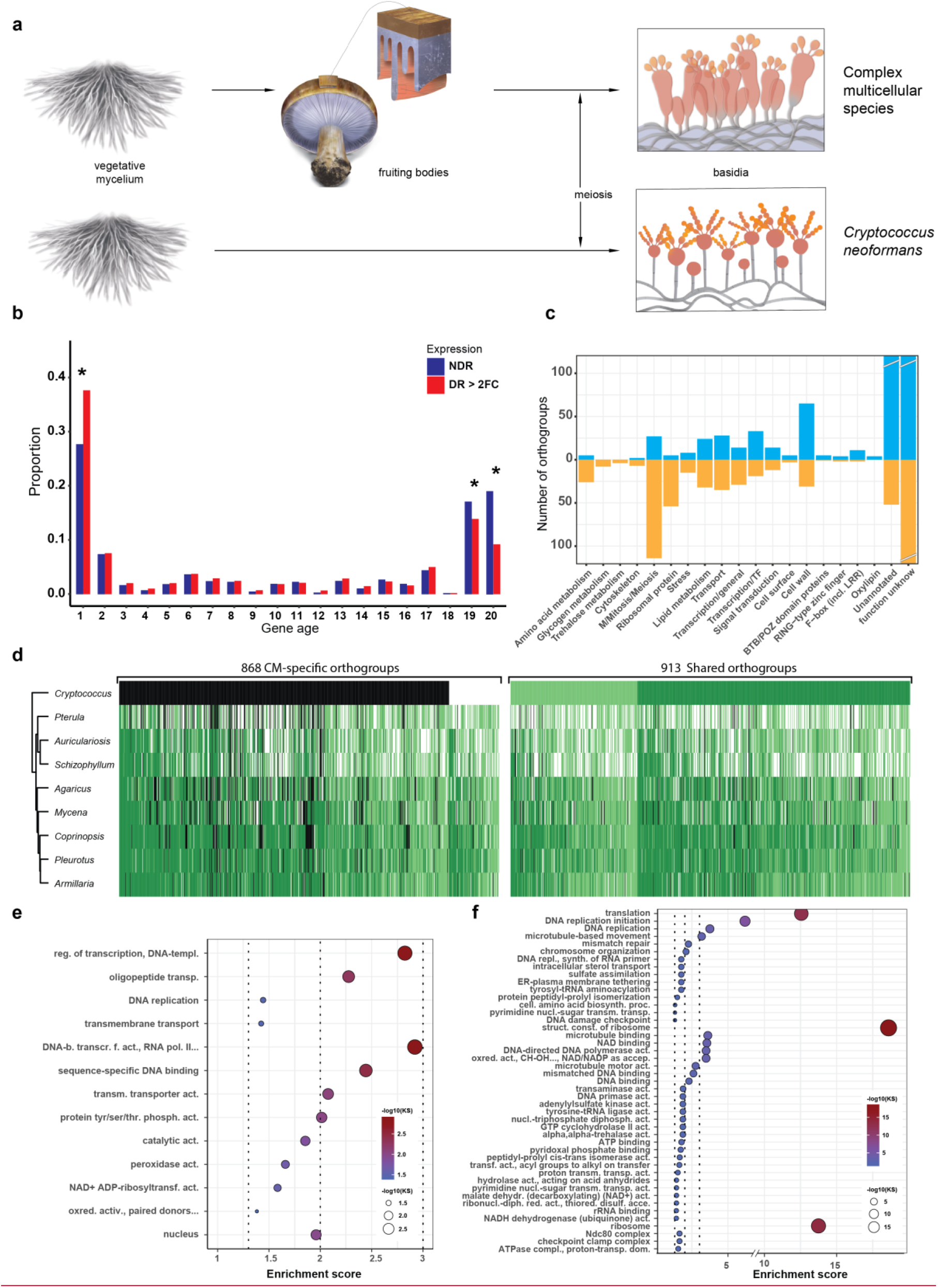
Conserved developmental expression in CM fungi. **a)** schematic representation of complex multicellular and simple development and the use of *C. neoformans* as a minimal model of sexual development in the Basidiomycota **b)** Enrichment analysis of developmentally regulated genes in different gene ages. **c)** Number of conserved orthogroups in different functional categories **d)** distribution of genes and their developmental expression in the nine species. Dark and light green refers to genes with developmental regulation at fold change > 4 and 2, respectively, whereas white and black denoted non-developmentally regulated and missing genes, respectively. Gene Ontology (GO) enrichment for **e)** CM-specific and **f)** Shared orthogroups. KS means the p-value of Kolmogorov-Smirnov test implemented in the R package ‘topGO’. On panels e and f, cutoff lines (dashed line) are drawn at enrichment scores corresponding to p=0.05, p=0.01 and p=0.001 (from left to right). GO terms are ordered by Kolmogorov-Smirnov p-values. See also Table S12 for GO enrichment details.

To distinguish genes related to basic sexual processes (sporulation, meiosis) from those related to CM, we reanalysed transcriptome data for sexual sporulation and basidium development of *Cryptococcus neoformans* (Liu et al., 2018). This species has a simple, non-CM development and we used it here as a minimal model of sexual development (Fig. 4/a/b). Of the 1,781 conserved orthogroups, 913 and 868 were developmentally expressed both in *C. neoformans* and CM species and only in CM species (Fig. 4/e), and are referred to as shared and CM-specific orthogroups, respectively. Of the 868 CM-specific orthogroups, 754 were completely missing in *C. neoformans* whereas 114 were present but not developmentally regulated (Fig 4/e, Table S8). Phylostratigraphic analyses showed that, on average, shared orthogroups contain evolutionarily old genes that predate the origin of Dikarya, while CM-specific orthogroups are younger (Fig. 5). Shared orthogroups included more housekeeping gene functions, such as mitosis/meiosis, general transcription factors or ribosomal proteins, whereas CM-specific orthogroups contained more genes encoding sequence specific transcription factors, cell wall remodelling, oxylipin biosynthesis, protein ubiquitination (F-box, BTB/POZ and RING zinc finger domain proteins) as well as functionally unclassified proteins (Fig. 4/d/f-g).

**Figure 5.**
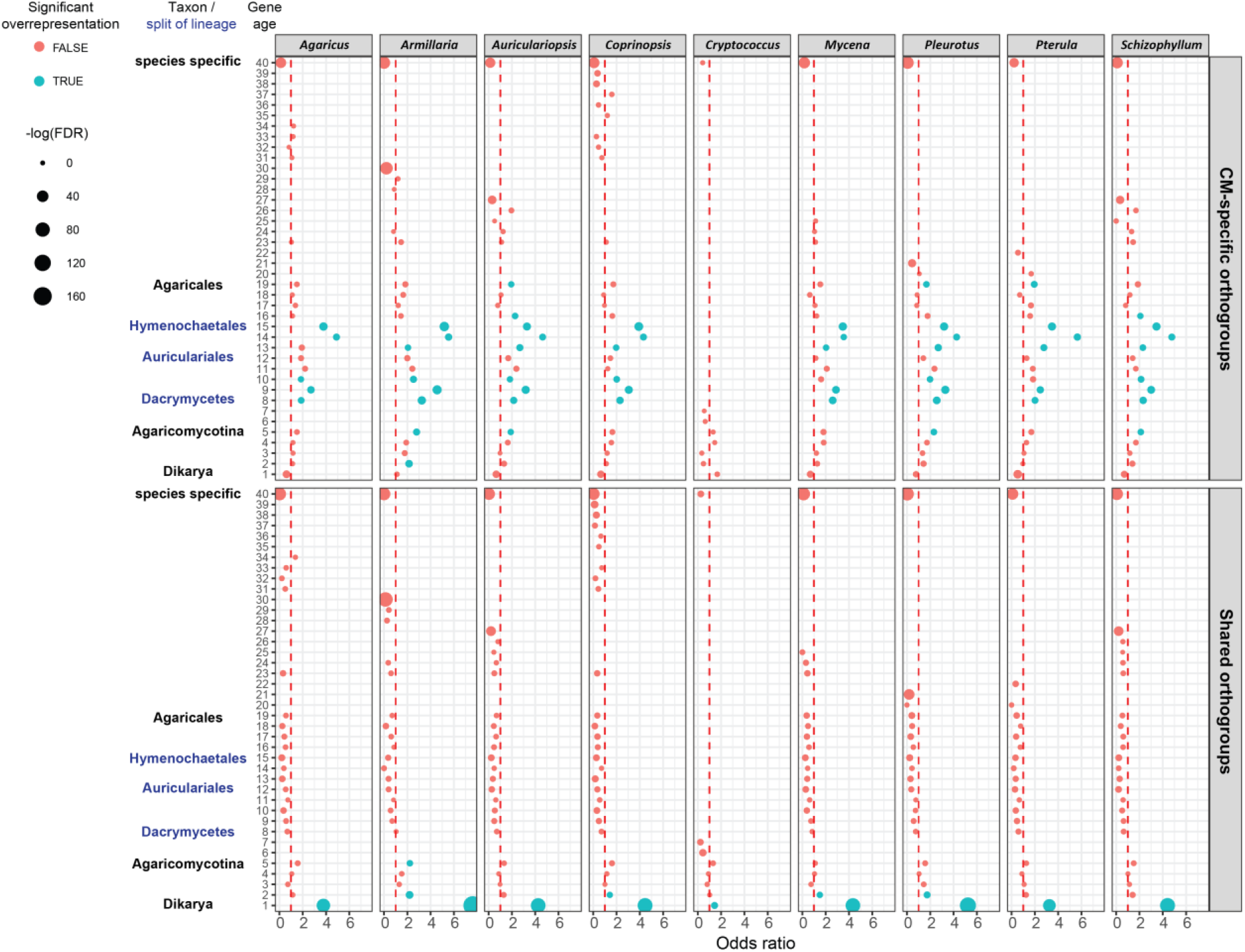
Gene age enrichment for shared and CM-specific conserved developmentally regulated orthogroups. CM-specific orthogroups are significantly enriched in the most recent common ancestors of lineages in which the first complex structures emerged. Y axis of the scatterplot represents relative gene age (see Fig. S29 for definition of ages). X-axis displays the odds ratio of Fisher’s exact test. If the odds ratio exceeds 1 (red dotted line) orthogroups are overrepresented in that gene age. Non-significant and significant (FDR p< 0.05) overrepresentation are marked with red and blue, respectively. The size of circles corresponds to the FDR corrected p-value.

Cell division related (DNA replication, meiosis, mitosis, DNA repair, etc.) and ribosomal protein encoding genes comprised the most frequent annotations in shared orthogroups (Fig 4/g). Meiosis happens in the basidia in both *C. neoformans* and CM fungi and associated genes showed clear peaks in their expression (Fig. S27). *C. neoformans* showed a single peak in meiotic/mitotic gene expression, whereas higher fungi like *P. ostreatus* showed two peaks, one corresponding to meiosis in gills and another to intense hyphal cell division (mitosis) in primordia. Ribosomal protein gene expression, as a proxy for the activity of protein synthesis, has been widely associated with cell growth and proliferation (Jorgensen et al., 2002; Kraakman et al., 1993). Ribosomal proteins showed an early peak in all species, while in CM species a second peak was also observed, coincident with meiosis and spore production in gills, possibly reflecting the requirement for increased protein synthesis (Fig. S28). We infer that the first ribosomal gene expression peak corresponds to an early, proliferative phase of development followed by the transition to growth by cell expansion (Krizsán et al., 2019), which gives the final shape and size of fruiting bodies before spore release.

Several cell surface proteins (fasciclins, ricin-B lectins and the PriA family) and putative cell wall remodelling enzymes (e.g. chitin- and glucan-active glycoside hydrolases, expansins, CE4 chitooligosaccharide deacetylases, laccases) previously linked to fruiting body morphogenesis (Pezzella et al., 2013; Xie et al., 2018) were shared between *C. neoformans* and CM species (Table S8), suggesting that these families are important for sexual morphogenesis in general, not restricted to CM. Cell wall remodeling enzymes have been hypothesised to produce fruiting body-specific cell wall architectures (Buser et al., 2010; Krizsán et al., 2019; Liu et al., 2021; Ohga, 2000); the upregulation of these in *C. neoformans* suggests a role during non-CM sexual processes as well, possibly in generating aerial hypha- or basidium-specific cell walls. Most genes related to glycogen metabolism also showed shared expression (Table S8). Glycogen has been known as a storage material in CM fruiting bodies, but our observations indicate that it may serve that role in *C. neoformans* too and possibly in sexual development in general. Notable transcription factors in share orthogroups included the light sensing white collar complex member WC-1, orthologs of *S. cerevisiae* sexual reproduction-related Ste12, a Basidiomycota-specific velvet factor as well as orthologs of the carbon catabolite repressor CreA (*A. nidulans*).

CM-specific orthogroups showed a phylostratigraphic enrichment in early mushroom-forming fungi (FDR<0.01, Fisher’s exact test, Fig. 5) and may correspond to innovations related to CM fruiting bodies, or, to a smaller extent, conserved genes lost in *C. neoformans* (e.g. hydrophobins and cerato-platanins). This observation complements our previous analysis (Krizsán et al., 2019) that could not resolve a clear signal of genetic innovation related to CM, possibly because of confounding effects of very conserved genes. In comparison to shared orthogroups, CM-specific orthogroups contained more transcription factors, genes related to cell wall biosynthesis/modification and defense (Table S9). Hydrophobins (probably the best-known fruiting body expressed genes) and cerato-platanins, as well as fatty acid desaturases and linoleate-diol synthases putatively related to biosynthesis of the signal molecule oxylipin were exclusively found in CM-specific orthogroups (Fig. 4/d). Hydrophobins and cerato-platanins are completely missing and defense-related genes are strongly underrepresented (1/4th that of CM species) in the genome of *C. neoformans*, probably as a consequence of the adaptation to a primarily yeast-like lifestyle (Nagy et al., 2014). The large number of conserved unannotated genes among CM-specific orthogroups underscore the still cryptic nature of CM development in fungi. Detailed descriptions of all conserved developmental genes will be given in a separate publication (Nagy et al in prep).

## Conclusions

In this study, we disentangled developmental transcriptomes of complex multicellular fungi, both in terms of mechanisms and function, using a comparative dataset that included the first developmental gene expression profiling data for *Pleurotus ostreatus* (oyster mushroom), the second most widely cultured fungus worldwide (Grimm and Wösten, 2018; Royse et al., 2017).

The availability of the genomes of both parental monokaryons (Alfaro et al., 2016; Riley et al., 2014) as well as new strand-specific RNA-Seq data allowed bioinformatic deconvolution of RNA editing, allele-specific expression and antisense transcription in the *P. ostreatus* transcriptome. We found virtually no evidence for RNA editing, whereas allele specific expression was abundant, which supports the only previously available report of ASE in CM fungi (Gehrmann et al., 2018). RNA editing has been recently reported in the Agaricomycetes (Wu et al., 2019, 2018; Zhu et al., 2014), however, in contrast to the Ascomycota (Lau et al., 2020; Liu et al., 2016; Teichert et al., 2017), it displayed no clear-cut enrichment of A-to-I compatible variants in three previous studies (Bian et al., 2019; Teichert, 2020; Zhu et al., 2014) or in this study. Rather, our final candidate RNA-editing sites merely alluded to potential polyA site- and/or read alignment inaccuracies leading us to conclude that RNA editing is not abundant in *P. ostreatus*. On the other hand, ASE was detected in thousands of genes in *P. ostreatus*, In *A. bisporus* ASE was interpreted as a regulated and adaptive mechanism that could, for example, aid the division of labour between nuclei (Gehrmann et al., 2018). We found that in *P. ostreatus* ASE is characteristic of young genes and likely arises from promoter divergence, which creates a cellular environment with divergent cis-regulatory alleles but identical trans-regulatory elements. Young genes are known to be under weaker evolutionary constraint than conserved ones, raising the possibility that ASE might arise neutrally in the transcriptome. This would be consistent with the neutral model of expression evolution (Fay and Wittkopp, 2008) and non-adaptive explanations, such as leaky regulation or transcriptional noise (Cheng et al., 2017; Khan et al., 2012; Shih and Fay, 2021; Wainer-Katsir and Linial, 2019). Under this interpretation, ASE may be a tolerated, rather than an adaptive phenomenon in CM fungi. However, even if neutral at the level of the individual, ASE may generate gene expression variation that can serve as substrate for adaptive evolution, similar to how transcription from random promoters can facilitate de novo gene birth (Van Oss and Carvunis, 2019). ASE may have important implications in mushroom breeding, where intraspecific hybrids harbouring cis-regulatory alleles with various levels of divergence may show differences in industrially relevant traits (Gehrmann et al., 2018).

Fruiting bodies integrate several distinct processes, some of which are conserved across all organisms (mitosis/meiosis), fungi (spore production), or specific to CM species (fruiting body morphogenesis). Analysing the evolutionary stratification of developmentally expressed genes helped us distinguish conserved genes related to basic sexual development from those putatively related to CM morphogenesis. As in the case of ASE, young genes might display more expression variance and noise across development, whereas genes with conserved developmental expression more likely provide clues about key CM functions. While our transcriptome-based approach has limitations as far as conservation of expression versus that of function is concerned, it proved relevant for understanding the evolution of CM. This may help for establishing a minimal model of sexual development (e.g. for pathogens like *C. neoformans*) in the Basidiomycota, that include several genes previously considered specific to fruiting bodies. Notable examples include fasciclins, which have been implicated in cell adhesion (Nagy et al., 2018), and the *PriA* family of secreted cell surface proteins (including *C. neoformans* cfl1 and dha1 Gyawali et al., 2017) with unknown function. Our strategy also identified regulatory genes which are developmentally expressed only in CM species, including certain transcription factors reported in mushrooms (e.g. hom1, fst3, fst4, wc1; Hou et al., 2020; Kamada et al., 2010; Ohm et al., 2011) but also novel ones, such as a velvet factor that is widely conserved in Agaricomycetes and showed stipe-specific expression in CM species.

Taken together, this study emphasises the complexity of transcriptome evolution in the transition to complex multicellularity in fungi. Our results suggest that CM transcriptomes are likely shaped by both adaptive and neutral processes and encourage non-adaptationist views of transcriptome evolution and analysis. We anticipate that the dissection of shared and CM-specific genes as well as allele-specific expression and natural antisense transcripts will help understanding the intricate gene regulatory networks governing development in both pathogenic and CM Basidiomycota and will contribute to the growth of mushroom developmental biology.

## Materials & Methods

### Growth condition, sampling and RNA-Sequencing

For fruiting the dikaryotic strain N001 (CECT-20600) of *Pleurotus ostreatus* (recently interpreted as *P. cf. floridanus* J. Li et al., 2019) we first prepared spawn by inoculating sterilized rye and incubating for 10 days. Pasteurized straw-based commercial oyster compost (95 vol%) and the colonized spawn (5 vol%) were mixed gently and three kilograms were filled into polyethylene bags. Bags were incubated in the dark at 27 °C and 85-90% relative humidity for 17 days. Next, bags were transferred to the growing room for fruiting at 18-19 °C, relative humidity 80-85% and 8/16h light/dark period (with approximately 1200 lux light intensity). We sampled vegetative mycelium (VM), six developmental stages and five tissue types, each in three biological replicates as explained in Fig. S1. VM was collected from the sawdust culture. In the case of stage 1 and 2 primordia (P1 and P2) the whole tissue was collected containing both stipe and cap initials. In the stage 3 primordium (P3) and the young fruiting body (YFB) stages, stipes and caps were sampled separately. We divided mature fruiting bodies (FB) into stipe (S), cap trama (C), cap cuticle (V). We defined cap (H) as the whole upper part of the fruiting body (in P3 and YFB) while cap trama (C) refers to just the inner part of cap without lamellae or cuticule (in FB). The last stage was the dedifferentiated cap trama (D), a dissection from inner cap tissue which was inoculated for 24h on a sterile PDA-agar plate, until the emergence of new hyphae. Tissue from 3-8 individual fruiting bodies was pooled for each replicate of each sample type.

*Pterula gracilis* CBS 309.79 was inoculated onto Malt Extract Agar plates with cellophane, and incubated at 25 °C for 25-27 days. For fruiting, plates were moved to a growth chamber at 15 °C under 10/14h light/dark period (light intensity: 11 μE m^−2^ s^−1^). VM samples were scraped off the cellophane after three days. We defined primordia (P) as small (<1mm) globose structures, young fruiting bodies as ~5 mm long awl-shaped structures, while structures longer than 10mm were considered mature fruiting bodies (Fig. S2).

Three biological replicates of each sample type were stored at −80 °C until RNA extraction. Tissue samples were homogenized with micropestles using liquid N2, and RNA was extracted by using the Quick-RNA Miniprep Kit (Zymo Research) according to the manufacturer’s instructions. Strand-specific cDNA libraries were constructed from poly(A)-captured RNA, using the Illumina TruSeq Stranded RNA-Seq library preparation kit, and sequenced on the Illumina HiSeq 4000/x platform in PE 2×150 format with 40 million reads per sample at OmegaBioservices (USA).

### Bioinformatic analyses of RNA-Seq data

New data for *P. ostreatus* and *Pt. gracilis* was reanalysed together with previously published transcriptomes of seven Basidiomycota species (Table S10). To remove adaptors, ambiguous nucleotides and any low quality read end parts, reads were trimmed using bbduk.sh and overlapping read pairs were merged with bbmerge.sh tools (part of BBMap/BBTools; http://sourceforge.net/projects/bbmap/) with the following parameters: qtrim=rl trimq=25 minlen=40. A two-pass STAR alignment (Veeneman et al., 2015) was performed against reference genomes with the same parameters as in our previous study (Krizsán et al., 2019) except that the maximal intron length was reduced to 3000 nt. Read count data was normalized using EdgeR (Robinson et al., 2010) as in our previous study (Krizsán et al., 2019). Expression levels were calculated as fragments per kilobase of transcript per million mapped reads (FPKM). Samples, such as FBCL and FBS of *Coprinopsis cinerea from* Krizsán et al. (2019), and stage 2 primordia (P2) of *P. ostreatus* were excluded from our analysis to avoid the signs of fruiting body autolysis and for quality reasons, respectively. Raw RNA-Seq reads have been deposited to NCBI’s GEO archive (GSE176181).

### Identification of developmentally regulated genes

Developmentally regulated genes were defined as in Krizsán et al. (2019). Genes with both two- and four-fold expression change were considered developmentally regulated and analyzed separately.

We checked if developmentally regulated genes physically clustered together and if they overlapped with predicted biosynthetic gene clusters in eight Basidiomycota species, including *P. ostreatus* and *Pt. gracilis* (Table S10). To this end, we first considered all developmentally regulated genes separated by six or less intervening genes to be clustered, and then calculated the probability of observing a cluster of that size using a binomial test, based on the expected number of developmentally regulated genes that would be found in the same sized gene window as the cluster if all developmentally regulated genes randomly distributed across the genome. Clusters with a probability of observation < 0.01 were designated ‘hotspots’. We then checked hotspots for overlap with predicted biosynthetic gene clusters using *de novo* antiSMASH v5 annotations (Medema et al., 2011). Conservation of hotspots between different fungal species was assessed by BLASTp (Altschul et al., 1990) using two metrics: percent gene content similarity, which is the number of genes in the query hotspot that are clustered together in the target genome; and percent FDBR similarity, which is the number of developmentally regulated genes in the query hotspot that are also developmentally regulated and clustered together in the target genome, using the same clustering criteria as above.

### Species tree and relative gene age estimation

Protein sequences of 109 whole genomes (Table S11) across Basidiomycota and Ascomycota (as outgroup) were downloaded from the JGI genome portal (Sep. 2019; Grigoriev et al., 2014; Nordberg et al., 2014). All-vs-all similarity search was carried out with MMseqs2 (Steinegger and Söding, 2017) using three iterations, and setting sensitivity to 5.7, max-seqs to 20,000, e-profile to 1e-4, a preliminary coverage cut-off to 0.2 and an e-value cut-off to 0.001. Then, an asymmetrical coverage filtering was performed where we required >=0.2 pairwise alignment coverage from the longer protein and >=0.8 from the shorter one, with the aim to omit aspecific partial hits while retaining gene fragments. Then Markov Chain Clustering with an inflation parameter 2.0 was performed using the ratio of ‘number of identical matches’ (Nident) and ‘query sequence length’ (qlen) as weight in the matrix. After clustering we removed contaminating proteins from gene families following the logic of Richter et al. (2018).

For species tree reconstruction we used 115 single copy gene families, which were present in >= 50% of the 109 species. Multiple sequence alignments were inferred using PRANK v.170427 (Löytynoja, 2014) and trimmed with TrimAL v.1.2 (-strict) (Capella-Gutierrez et al., 2009). Trimmed MSA-s shorter than 100 amino acid (AA) residues were discarded. Best partitioning schemes, substitution models and species tree reconstruction were performed under Maximum Likelihood (ML) in IQ-TREE v1.6.12 (Minh et al., 2020).

For gene tree reconstructions, gene families which contained at least four proteins were aligned with MAFFT LINSI v7.313 (Katoh and Standley, 2013) algorithm or with FAMSA v1.5.12 (Deorowicz et al., 2016) and trimmed with TrimAL (gt-0.4). We inferred gene trees for each of the alignments in RAxmlHPC-PTHREADS-AVX2 under the PROTGAMMAWAG model of sequence evolution and assessed branch robustness using the SH-like support (Stamatakis, 2014). Rooting and gene tree/species tree reconciliation were performed with NOTUNG v2.9. (Darby et al., 2016) using an edge-weight threshold of 90. Then, a modified version of COMPARE (Nagy et al., 2014) was used to delineate orthogroups within gene trees.

Orthogroups were used to define relative gene ages (hereafter: gene age) as the species tree node to which the Most Recent Common Ancestor (MRCA) of species represented in the orthogroup mapped. Enrichment of gene sets in gene age categories were analysed with Fisher’s exact test (R core team 2020).

### Transcriptome age index

Transcriptome age index for each developmental stage of the nine investigated species was computed as described previously (Domazet-Lošo and Tautz, 2010) with slight modifications, using the following formula: 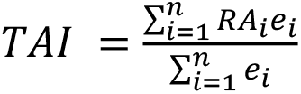, where RA_i_ represents the relative age of gene *i, e*_*i*_ is the log2 FPKM value of gene *i* at the given stage and *n* is the total number of genes. If available, tissue specific expression values were averaged for each developmental stage. The TAI values of the investigated developmental stages were computed for each replicate, then averaged.

### Orthology based on reciprocal best hits

To characterize the conservation of developmental genes, we defined single copy orthologs from the nine species based on reciprocal best hits between proteins. This strategy was stricter than the above-mentioned orthogroup definition, and was required to obtain functionally highly similar protein sets for comparing developmentally regulated genes. Proteins of each species were searched against the proteomes of other eight species using the RBH module of MMSeqs2 with an e-value cut-off of 1e-5. To remove spurious reciprocal best hits, we excluded a protein from the RBH group if its bit score was at least three times lower than the mean bit score of other hits of that query (self-hit excluded) and it shared <50 % of its hits with those of the query. The orthogroups obtained this way comprised considerably more focused gene sets than the approach used in Krizsán et al. (2019).

### Annotation of genes, gene families

We detected conserved domains in proteins using InterProScan-5.47-82.0 (Jones et al., 2014). Enrichment analysis on IPR domains was performed with Fisher’s exact test (R Core Team, 2020), while enrichment analysis on gene ontology categories was carried out using the R package topGO 2.44.0 (Alexa and Rahnenfuhrer, 2020). Proteins were further characterized by the best bidirectional hits to proteins of the model organisms *Saccharomyces cerevisiae* (Engel et al., 2014), *Schizosaccharomyces pombe* (Wood et al., 2012), *Neurospora crassa* (Galagan et al., 2003) and *Aspergillus nidulans* (Cerqueira et al., 2014).

### Identification of Natural Antisense Transcripts

Natural antisense transcripts (NAT) were defined as *de novo* assembled transcripts located antisense to a gene >200 nucleotide (nt) long, not showing similarity to structural RNA species, not overlapping with UTRs of neighbouring genes, and showing an expression above a given cut-off (Fig. S18). For *de novo* transcript assembly, quality filtered reads were first mapped to the reference genome using STAR_2.6.1a_08-27 (Veeneman et al., 2015). After identifying splice sites, a second mapping was performed. StringTie version 2.0.3 (Pertea et al., 2015) was used to generate a genome-guided *de novo* transcriptome assembly and annotation. Transcripts shorter than 200 nt or supported by <5 reads in a single sample were excluded. Output GTF files from each sample were merged, compared to the reference annotation with gffcompare v0.11.2 (Pertea and Pertea, 2020) and transcripts with exonic overlap on the opposite strand (i.e. antisense; class_code x) were retained. To exclude transcripts mapped to repeats, RepeatModeler v2.0 (Flynn et al., 2020) was used to identify repeat regions of genomes. Conserved structural RNA transcripts (tRNAs, U2 spliceosome, ribosomal RNAs, Hammerhead ribozymes) were identified with Infernal 1.1.3 (Nawrocki and Eddy, 2013) based on the Rfam database (Kalvari et al., 2021) and were removed. Fungal genomes are densely packed with genes, raising the possibility that a detected transcript is actually the UTR region of the closest gene (Rhind et al., 2011). To avoid identifying UTR regions as NATs, candidate NATs showing a strongly correlated expression (Pearson’s P-value < 0.05) and located <500 nt from the closest coding genes on the same strand were discarded from further analysis. Coding potential of NATs were characterized based on the default cut-off of the Coding Potential Calculator CPC2 (Kang et al., 2017).

Expression level of NATs was quantified with the FeatureCounts R package (Liao et al., 2014) based on the union of the exons per transcript. FPKM calculations were carried out as mentioned above, only transcripts with at least five mapped reads in at least three samples were retained. Developmental regulation of potential NATs was calculated as for genes. To assess conservation of NATs we mapped the transcripts on the genomes of the 109 species with minimap2 v 2.17 (options: -k15 -w5 --splice -g2000 -G200k -A1 -B1 -O1,20 -E1,0 -C9 -z500 -ub --junc-bonus=9 --splice-flank=yes) (Li, 2018).

### RNA editing and allele specific expression

To estimate the importance of RNA editing and Allele Specific Expression (ASE) during fruiting body formation of *P. ostreatus* we evaluated mismatches in Illumina reads according to their potential origin (RNA-editing, ASE, noise). A custom pipeline (Fig. S12) was constructed to first classify mismatches either as candidates for “RNA editing” or “allele-specific”. Then, these mismatches were analysed further in more specialized pipelines. First, we hard trimmed 10-10 nucleotides from both the 3’ and 5’ end of already quality trimmed reads to decrease the impact of sequencing errors during variant calling. A two-round STAR alignment was performed against both parental genomes (PC15 and PC9) as references, with the above mentioned parameters. Variants were identified with a custom awk script excluding bases with a Phred quality value below 30. Nucleotides differing the same way from both parental alleles were considered technical errors (caused by PCR amplification, sequencing or alignment), somatic mutations or RNA editing. Therefore, such mismatches were transferred to the RNA editing specific pipeline. In contrast, variants which differed only from one of the parental genomes were attributed to allele specific expression. The first part of the pipeline yielded lists of variants which were further analysed either in the RNA editing specific pipeline or in allele specific expression pipeline, as follows.

In the RNA editing pipeline (Fig. S12) we re-called variants for sites with pysamstats 1.0.1 (https://github.com/alimanfoo/pysamstats; min-base quality 30 and max depth to 500,000) that were input to the RNA editing pipeline. We continued the analysis with only those variants which had >=3 supporting reads, mapped to both reference genomes and the total read support did not differ more than five times between the two parental mappings, in order to avoid signal coming purely from the differential mapping to the two reference genomes (e.g. erroneous alignment). Further, we removed variants where read coverage was <10, a single variant was supported by <3 reads, or the proportion of the variant was below 0.1% in order to reduce the effect of technical errors, but retain editing sites. Because erroneous alignment around splice sites can produce variants indistinguishable from editing events, we discarded variants in which multiple sites with mismatches grouped within 3 nt distance of each other and in which the proportion of gaps exceeded 80% of the read coverage. After this step we kept only variants which were present in at least two biological replicates. In addition, for a variant to be considered an RNA editing site, it had to be significantly more frequent across all samples (Wilcoxon rank sum test with p<0.01) than any other nucleotide at that site (except reference). Finally, we considered a site an RNA editing site, if the geometric mean of its frequencies across the three replicates exceeded 1%. Relative to the editing site, −3 upstream and +3 downstream surrounding sequences were extracted with rtracklayer package (Lawrence et al., 2009) and motifs were searched with the seqlogo package (Bembom and Ivanek, 2020).

In the allele specific expression pipeline (Fig. S12), only previously assigned candidate allele specific SNPs were considered. All reads were assigned to the parental genome to which it exhibited a smaller Hamming distance (Hd=number of SNPs). We assigned a read as indecisive if i) Hd>1 from both reference genomes ii) Hd>15 from any of the reference genomes (too divergent read) or, iii) if the Hd was equal to both parental genomes. FPKM values were calculated as described above.

To describe the relative expression between the two parental nuclei (AS ratio) the number of PC15 reads was divided by the sum of parental specific reads (PC15 + PC9) for each gene (g) in each sample (s): 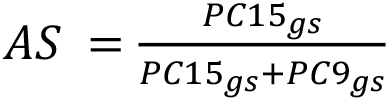. An AS ratio close to 1 means dominant expression from the PC15 nucleus, whereas an AS ratio close to 0 means dominant expression from the PC9 nucleus. An AS ratio ~0.5 indicates equal expression from both nuclei. AS ratios were considered equal (set to 0.5) if i) the expression was too low (FPKM<2), ii) the number of decisive reads was <16, iii) the proportion of indecisive reads was greater than 80%. We calculated two further measures, Chromosome Read Ratio (CRR) and Nuclear Read Ratio (NRR) introduced by Gehrmann et al., (2018), which represent the FPKM values of PC15 nucleus divided by the FPKM values of PC9 summed over chromosomes, and over all genes, respectively.

We identified genes with two- (S2) and four-fold (S4) shifted expression between the two nuclei at AS cut-off values of AS<0.31 or AS>0.68 (corresponding to 5% quantile of all AS ratio values) and AS<0.2 or AS>0.8, respectively. For passing through these filters and geometric means of replicates had to reach the upper limits (0.68 or 0.8) for PC15 specific ASE or less than lower limits (0.31 or 0.2) for PC9 specific ASE.

In order to understand how ASE may arise mechanistically, we compared putative promoters of ASE genes among parents, defined as the region from spanning 1000 nt upstream to 200 nt downstream of the transcription start site. Differences between the two parental regions were expressed as percent identity. For this analysis the meanwhile released, improved version of PC9 was used (Lee et al., 2021). Protein sequences of parents were aligned with PRANK and a ML distance was calculated under WAG model (dist.ml function from phangorn R package Schliep, 2011). Gene pairs with putative promoters <75% similar or where protein ML distances were >0.5 and alignment coverage <0.7 were removed to avoid potential orthology assignment errors. To identify the strength of selection for these genes we inferred ω (dN/dS ratios) under two evolutionary models using CodeML, a program from PAML 4.4 (Yang, 2007). For this, CDSs from the genomes of species in the Pluteoid clade (Table S11) were extracted using GenomicFeatures packages (Lawrence et al., 2013). 1:1 orthologs were detected with MMSeqs RBH function and codon aligned using –code option of PRANK. Reference tree for CodeML was extracted from the species tree, and ω values were calculated under one-ratio (M0) model assuming that ω has been constant throughout the tree and free‐ratio model (fb) allowing an independent ω for each branch in the tree. For statistical comparisons the Kruskal-Wallis rank-sum test with Nemenyi post hoc test or paired Wilcoxon signed-rank test and Fisher’s exact test were implemented in R (R Core Team, 2020).

## Acknowledgements

The authors acknowledge support by the Hungarian National Research, Development, and Innovation Office (contract No. GINOP-2.3.2-15-2016-00052), the “Momentum” program of the Hungarian Academy of Sciences (contract No. LP2019-13/2019 to L.G.N.) and the European Research Council (grant no. 758161 to L.G.N.).

## Data Availability Statement

The RNA-Seq data was deposited in the NCBI’s Gene Expression Omnibus (GEO) Archive at www.ncbi.nlm.nih.gov/geo (accession no. GSE176181). Other data used in this study are available on request from the corresponding author.

## Supplementary Figures for

**Figure S1.**
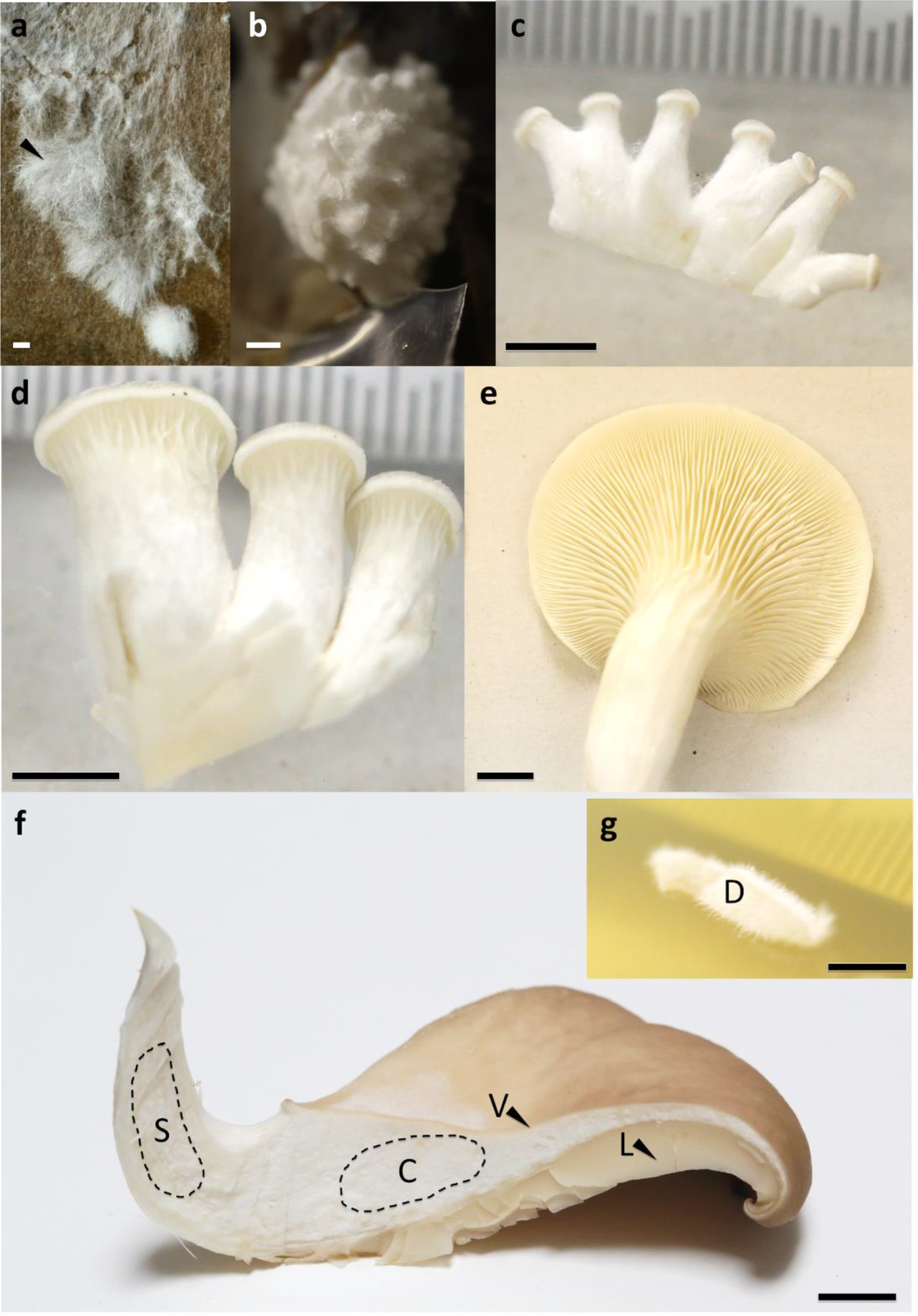
Sampled developmental stages and tissue types during fruiting body formation of *Pleurotus ostreatus*. a) ‘VM’ vegetative mycelium b) ‘P1’ stage 1 primordium; c) ‘P3’ stage 3 primordium; d) ‘YFB’ young fruiting body; e) ‘FB’ fruiting body; f) dashed areas and black arrows show different tissue types ‘S’ stipe, ‘C’ cap trama, ‘D’ dedifferentiated tissue of cap trama, ‘L’ lamellae, ‘V’ cuticle. Bars represent 1 mm in a and b, 5 mm in c and d while 1 cm in e-g.

**Figure S2.**
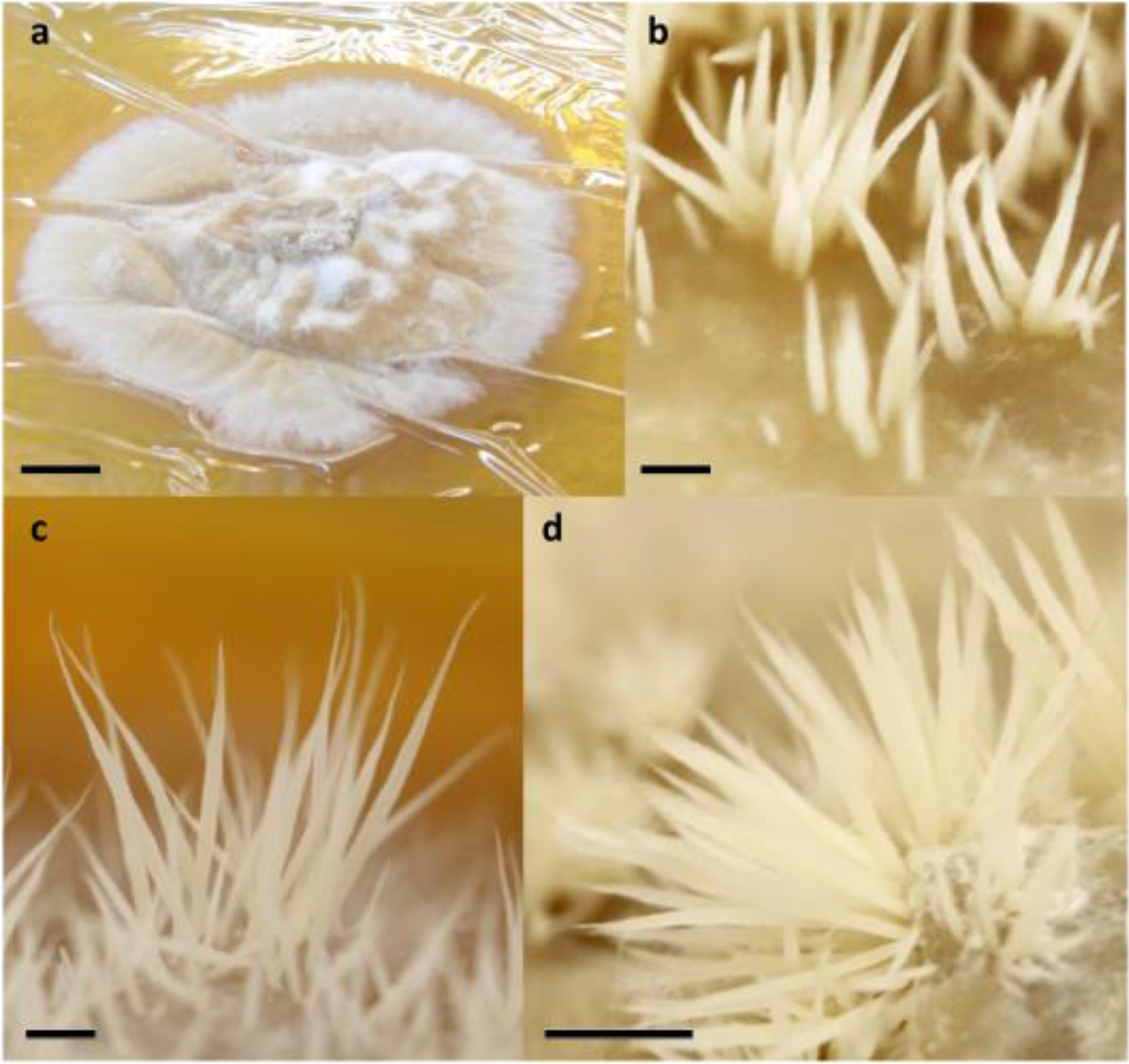
Sampled developmental stages during fruiting body formation of *Pterula gracilis*. a) ‘VM’ vegetative mycelium; b) ‘P’ primordium; c) ‘YFB’ young fruiting body; e) ‘FB’ fruiting body; Bars represent 1 mm in b-c 5mm in d while 1 cm in a.

**Figure S3:**
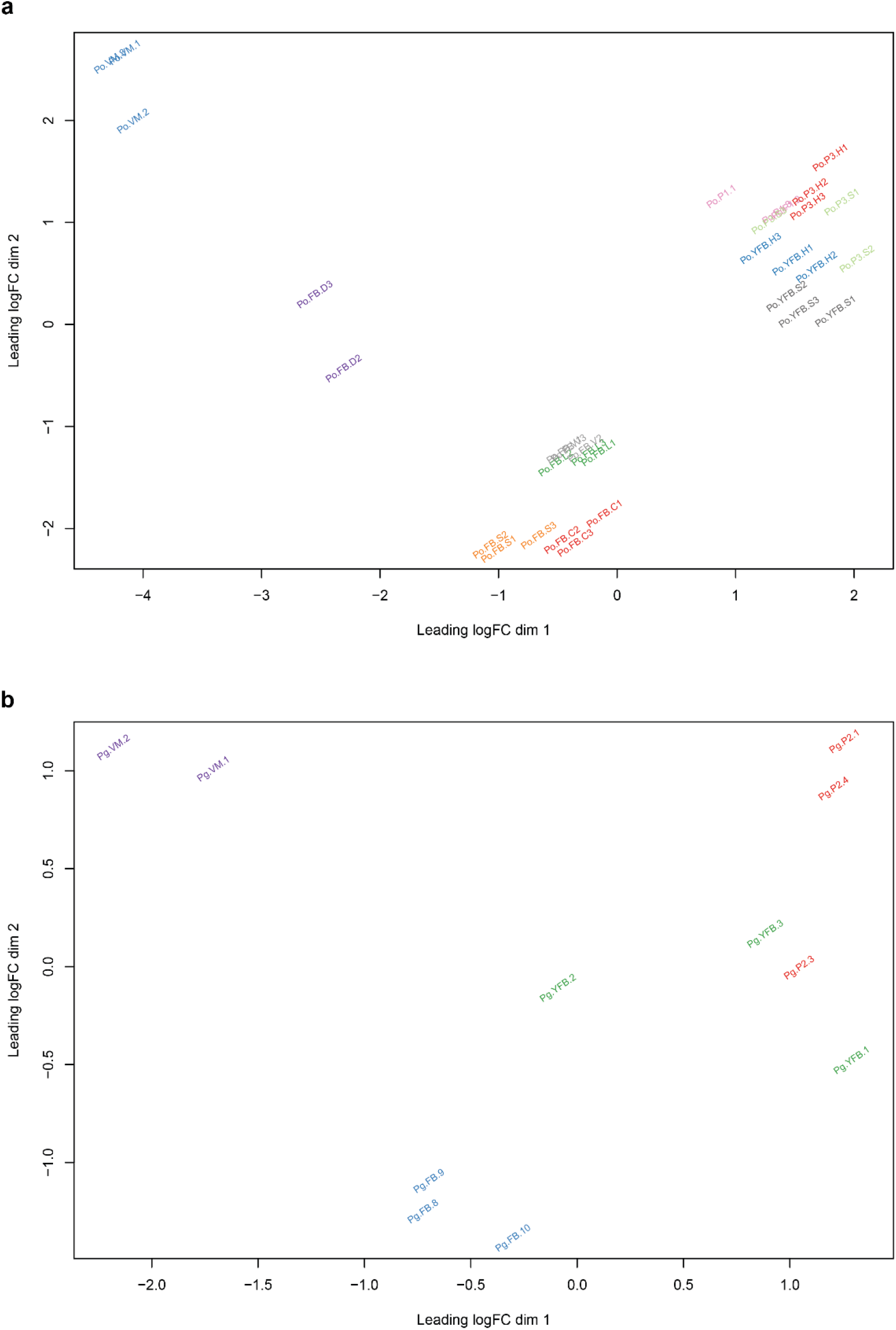
MDS plot based on the expression of genes in a) *Pleurotus ostreatus* and b) *Pterula gracilis*. Abbreviations as follows: ‘VM’ vegetative mycelium; ‘P1’ stage 1 primordium; ‘P3’ stage3 primordium; ‘YFB’ young fruiting body, ‘FB’ fruiting body; ‘H’ cap (entire); ‘C’ cap trama (only the inner part, without Lamellae, or skin); ‘L’ lamellae; ‘S’ stipe; ‘V’ cuticle; ‘D’ dedifferentiated tissue of cap.

**Figure S4:**
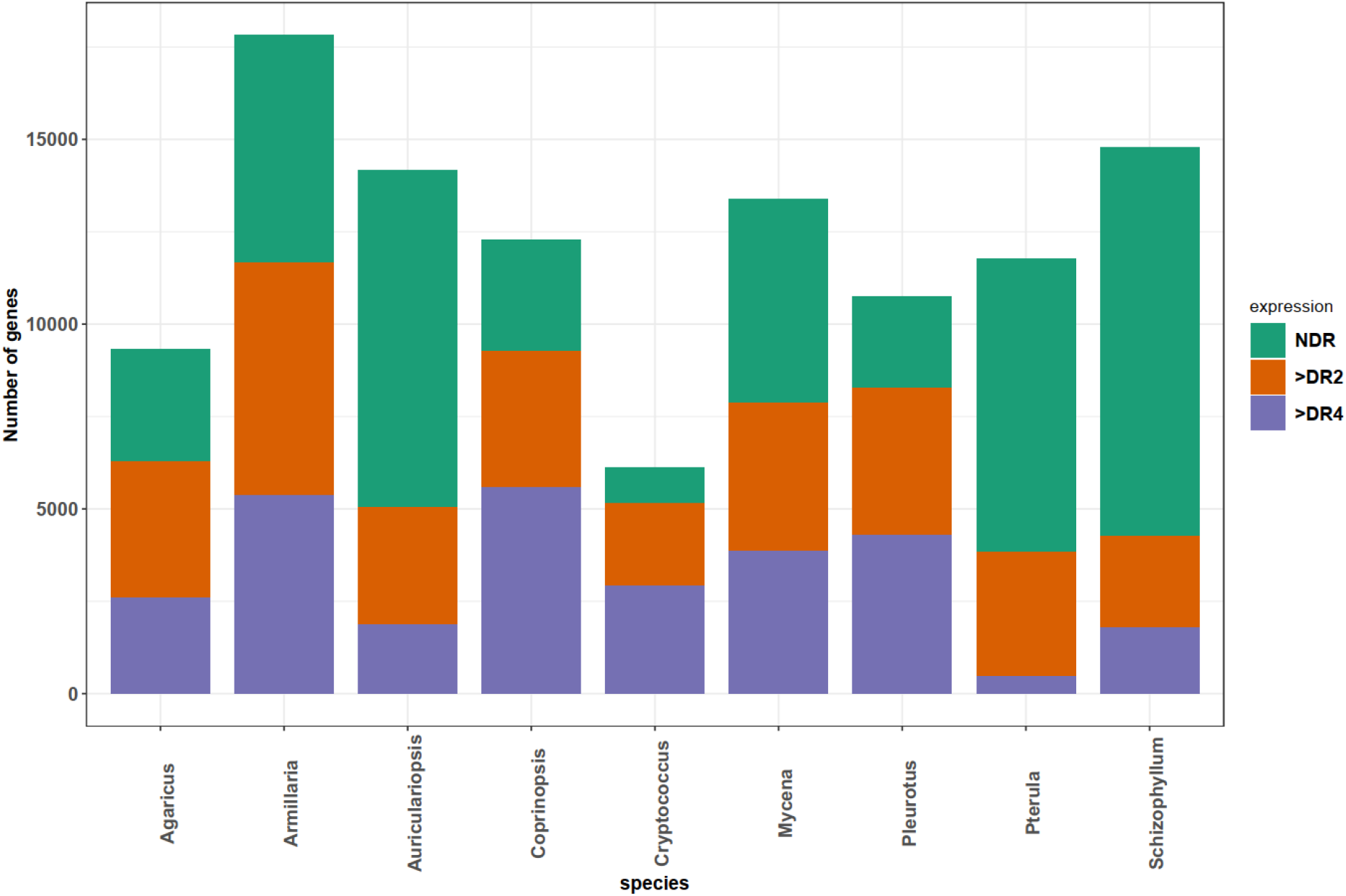
The distribution of developmentally regulated genes in each species. Abbreviations: NDR not developmentally regulated; >DR2 Developmentally regulated with at least 2 Fold Change; >DR4 Developmentally regulated with at least 4 Fold Change.

**Figure S5:**
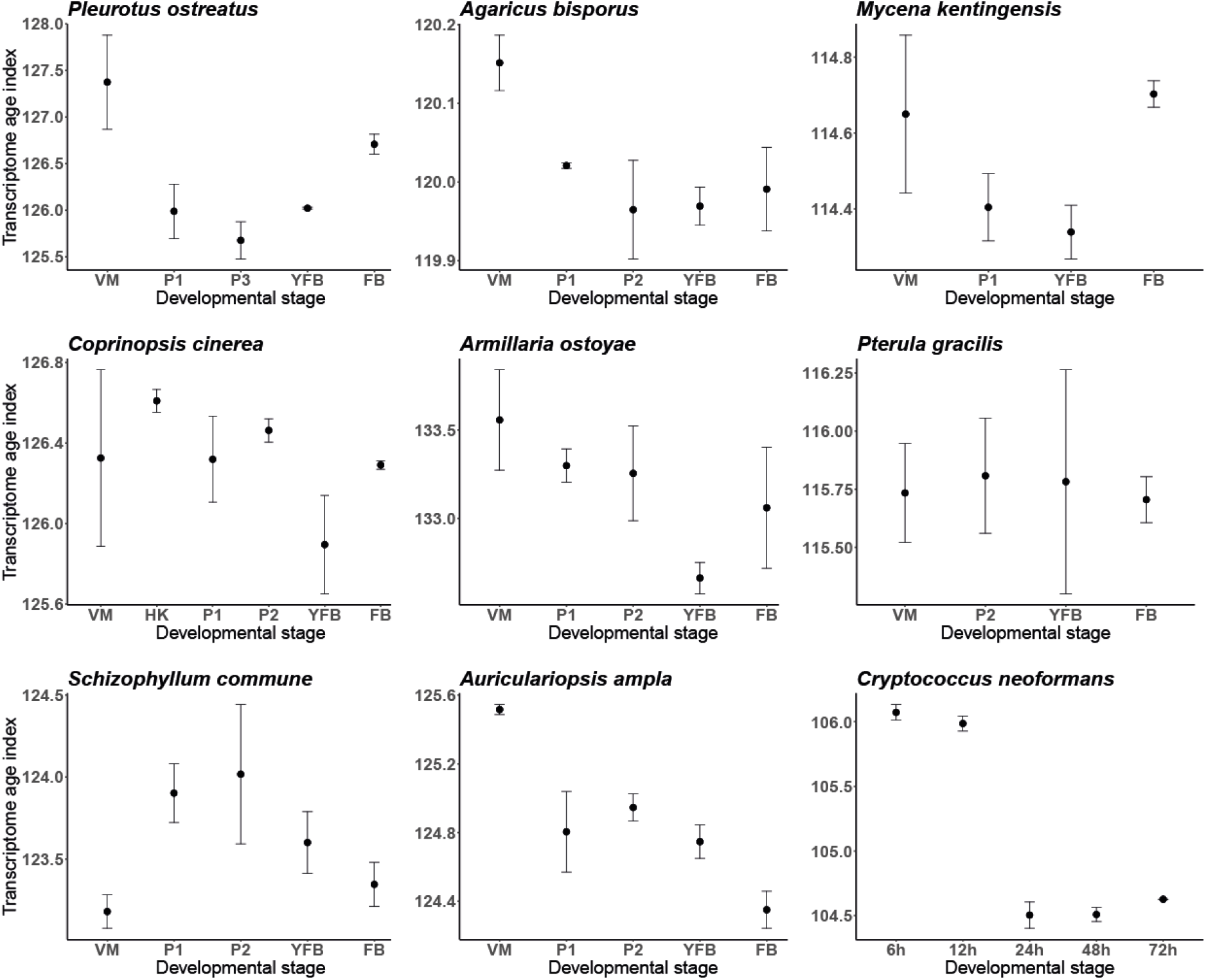
Transcriptome age index of the different developmental stages of the nine studied fungal species. The mean values of the transcriptome age indexes from three replicates were plotted throughout the development of eight fruiting body forming fungi and through the process of the basidium differentiation of *C. neoformans*. Lower TAI values represent older transcriptomes. We observed an hourglass-like pattern in case of *P. ostreatus*, *M. kentingensis*, *A. bisporus*, with stage 3 primordia, young fruiting body, and stage 2 primordia having the lowest TAI values, respectively. No consistent pattern was observed in the remaining six species. Abbreviations: VM: vegetative mycelia, HK: hyphal knot, P1: stage 1 primordia, P2: stage 2 primordia, P3: stage 3 primordia, YFB: young fruiting body, FB: fruiting body. Transcriptome Age Index (TAI) for the nine species.

**Figure S6:**
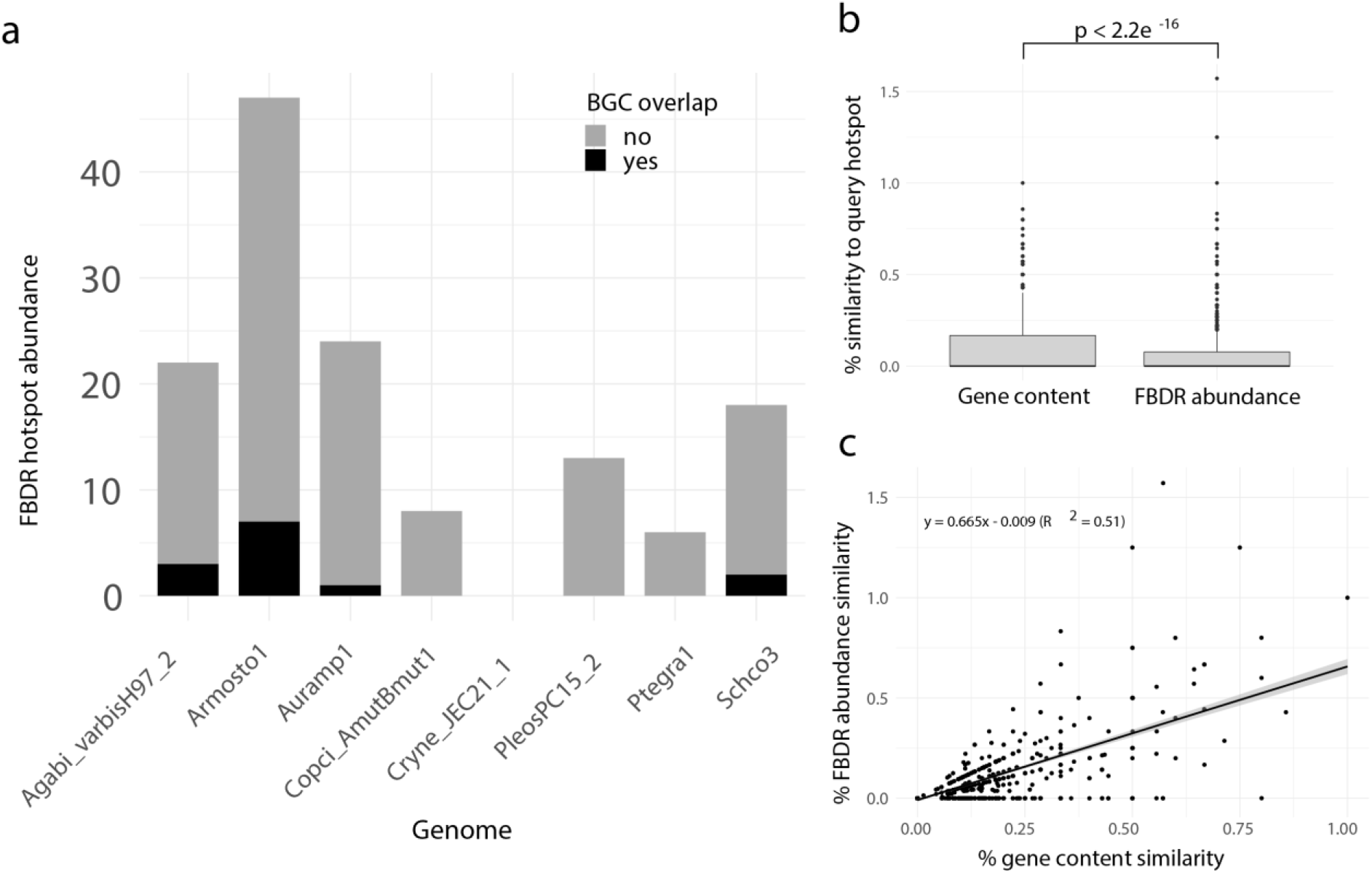
Developmentally regulated (FBDR) genes occasionally cluster together in genomic ‘hotspots’. a) A bar chart summarizing the number of FBDR hotspots detected per genome (total = 153), and the degree of overlap between FBDR hotspots and predicted biosynthetic gene clusters (BGCs). b) A box-and-whisker plot summarizing the distribution of % gene content conservation and the distribution of % FBDR abundance conservation of all 153 hotspots when searched for in genomes other than the one in which they are found. Significance between distributions determined by the Wilcoxon Rank Sum test. c) Scatterplot and linear regression describing relationship between the % gene content conservation and % FBDR abundance conservation of each of the 153 hotspots when searched for in genomes other than the one in which they are found (number of observations = 153 hotspot queries x 8 target genomes). % FBDR abundance conservation may exceed 100% if more FBDR genes are found in target regions as compared with query hotspot. Shaded region around the fitted regression line represents that 95% confidence interval.

**Figure S7:**
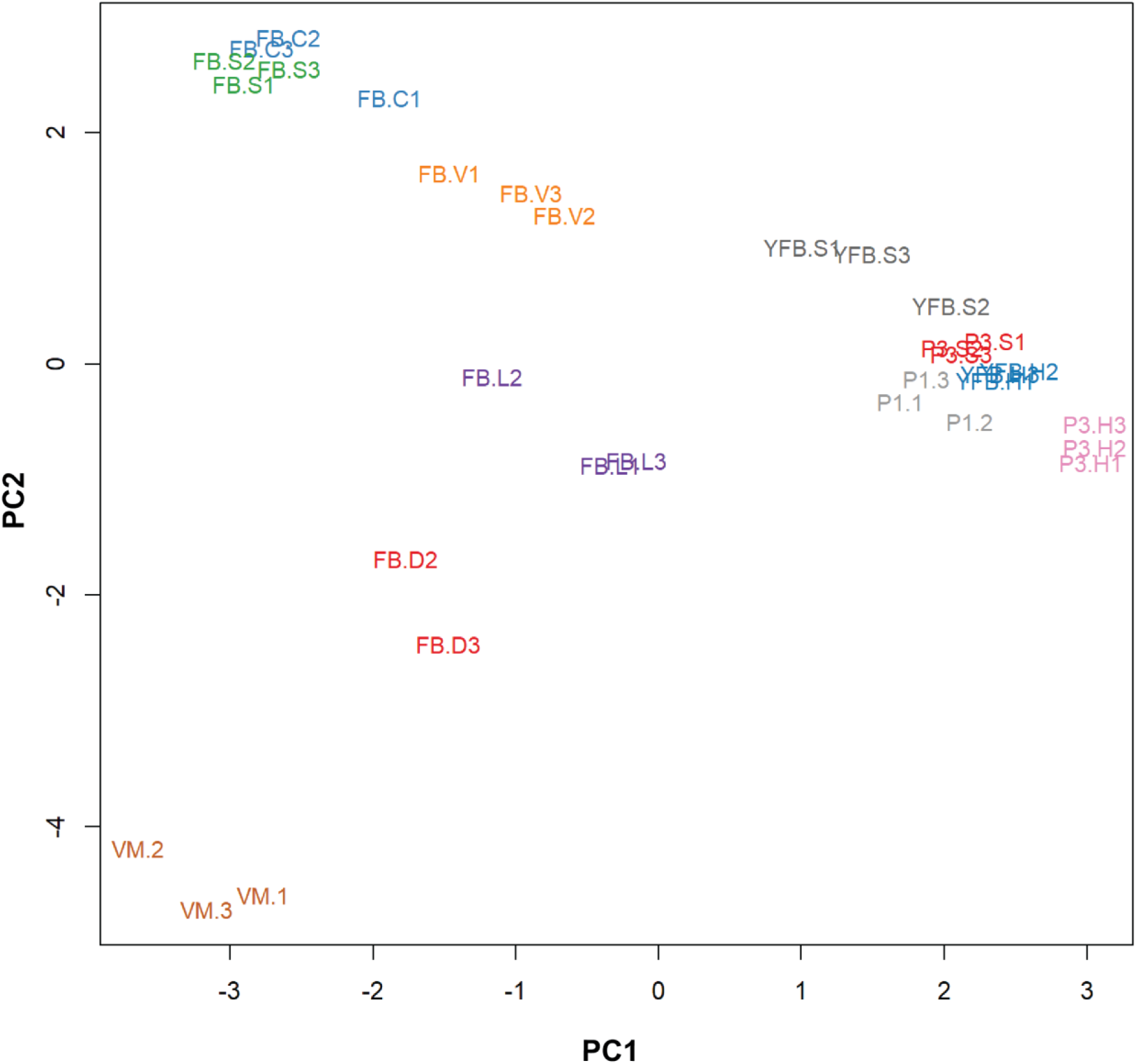
Principal Component Analysis based on AS ratio. Replicates of stages clustered together and therefore Allele Specific Expression showed a development- and tissue-specific pattern. Abbreviations as follows: ‘VM’ vegetative mycelium; ‘P1’ stage 1 primordium; ‘P3’ stage3 primordium; ‘YFB’ young fruiting body, ‘FB’ fruiting body; ‘H’ cap (entire); ‘C’ cap trama (only the inner part, without Lamellae, or skin); ‘L’ lamellae; ‘S’ stipe; ‘V’ cuticle; ‘D’ dedifferentiated tissue of cap.

**Figure S8:**
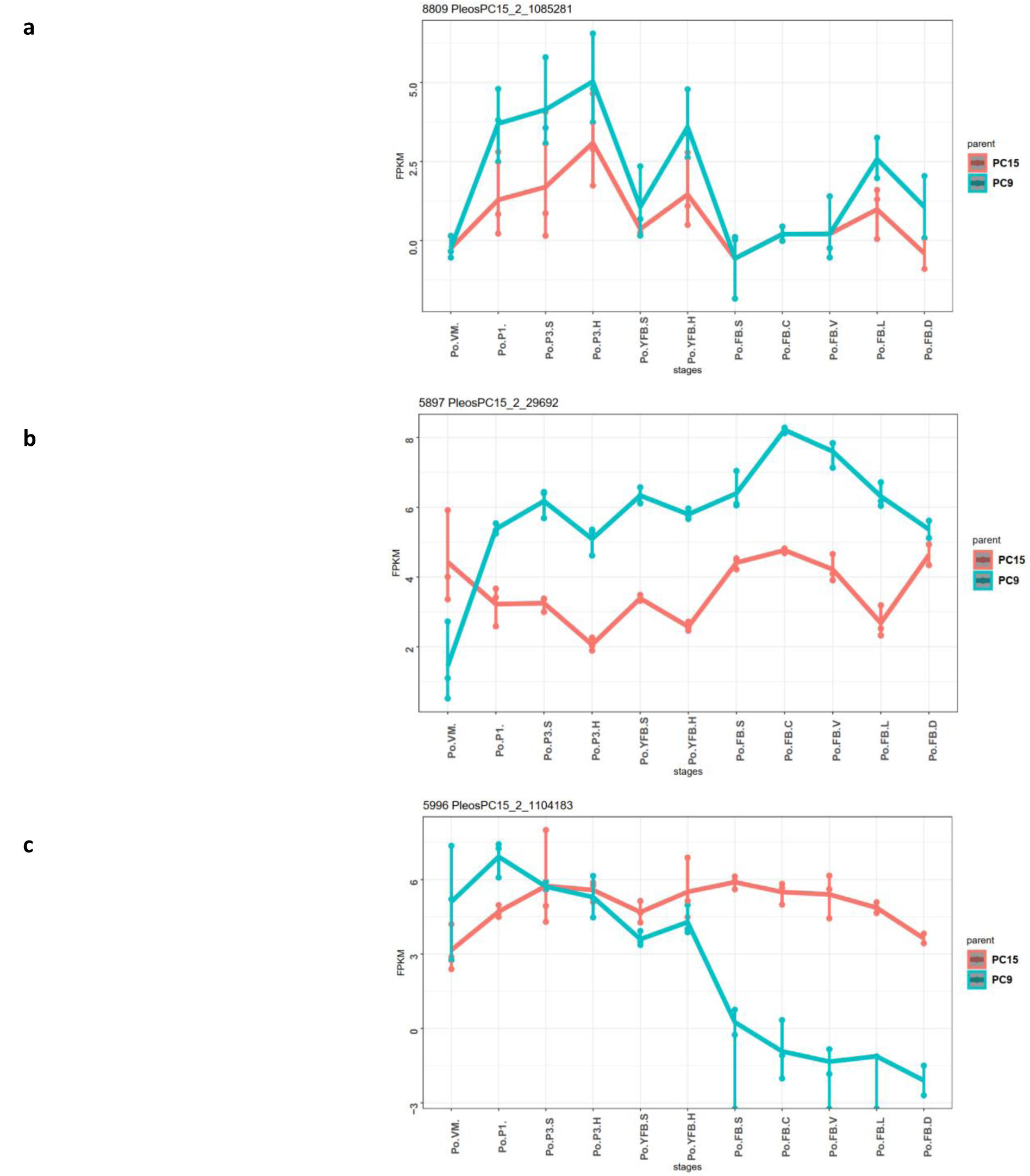
Examples for Allele Specific Expression (ASE) patterns during fruiting body formation of *Pleurotus ostreatus*. Gene expressions (log2 transformed FPKM) from the two nuclei are colored with blue (PC9) and red (PC15). *P. ostreatus* gene- and protein-IDs (PleosPC15_2_) are displayed in each plot as a title **a)** a conserved CON6-family sporulation-specific gene. **b)** Mannose-6-phosphate isomerase (IPR016305) **c)** a gene encoding a protein with F-box domain (IPR001810) and Leucine-rich repeat (IPR032675), where expression from PC9 is ‘turned off’ in mature fruiting bodies. Abbreviations as follows: ‘VM’ vegetative mycelium; ‘P1’ stage 1 primordium; ‘P3’ stage3 primordium; ‘YFB’ young fruiting body, ‘FB’ fruiting body; ‘H’ cap (entire); ‘C’ cap trama (only the inner part, without Lamellae, or skin); ‘L’ lamellae; ‘S’ stipe; ‘V’ cuticle; ‘D’ dedifferentiated tissue of cap.

**Figure S9:**
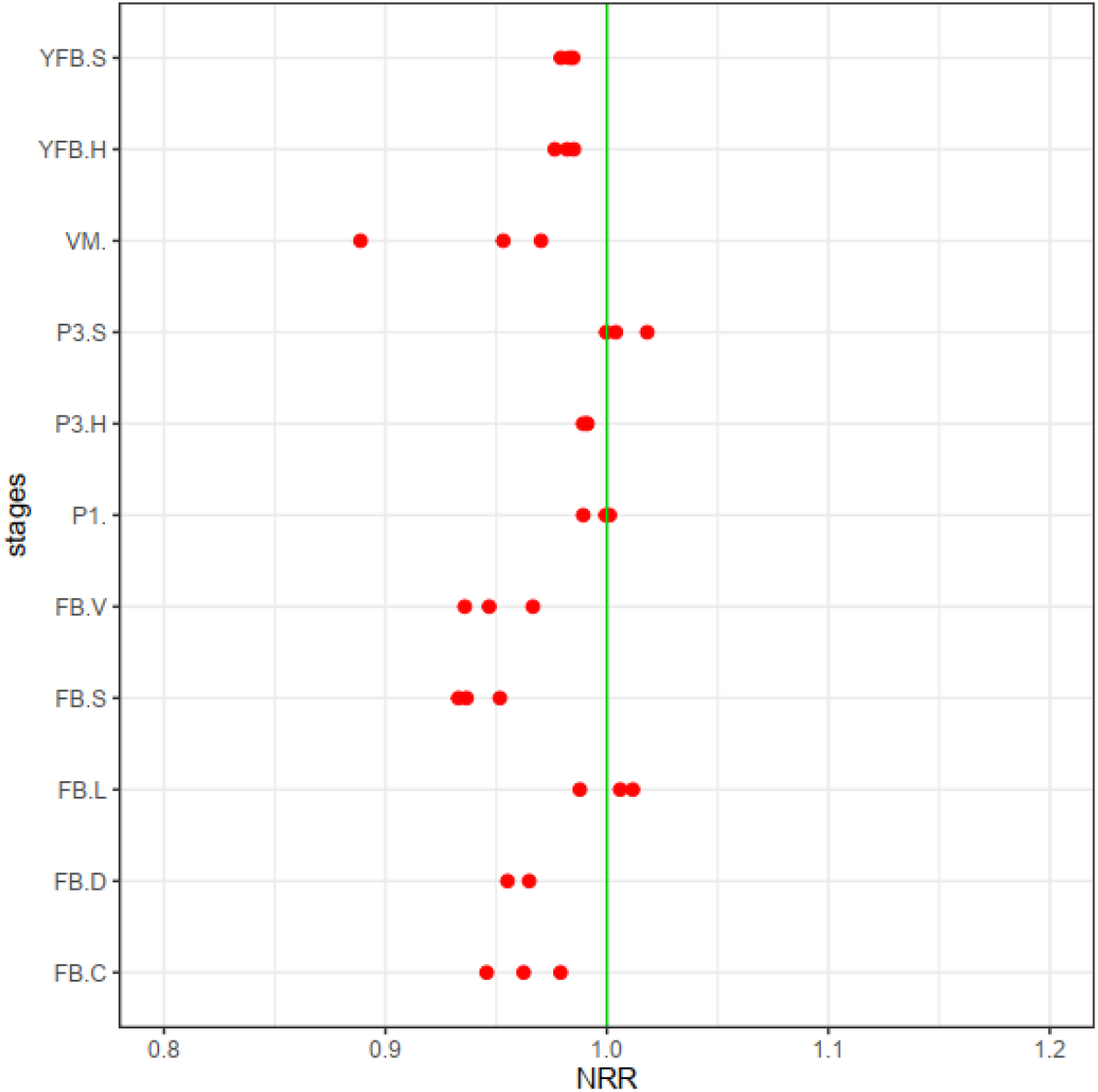
The Nuclear Read Ratio (NRR, the contribution of nuclei to the total expression) across stages and tissues. Red dots represent the biological replicates separately. A slight dominance of PC9 was observable (mean NRR across samples 0.97; max difference 12.5% Values smaller than 1 mean the dominance of the PC9 nucleus over PC15 while the opposite marks the dominance of PC15. Abbreviations as follows: ‘VM’ vegetative mycelium; ‘P1’ stage 1 primordium; ‘P3’ stage3 primordium; ‘YFB’ young fruiting body, ‘FB’ fruiting body; ‘H’ cap (entire); ‘C’ cap trama (only the inner part, without Lamellae, or skin); ‘L’ lamellae; ‘S’ stipe; ‘V’ cuticle; ‘D’ dedifferentiated tissue of cap.

**Figure S10:**
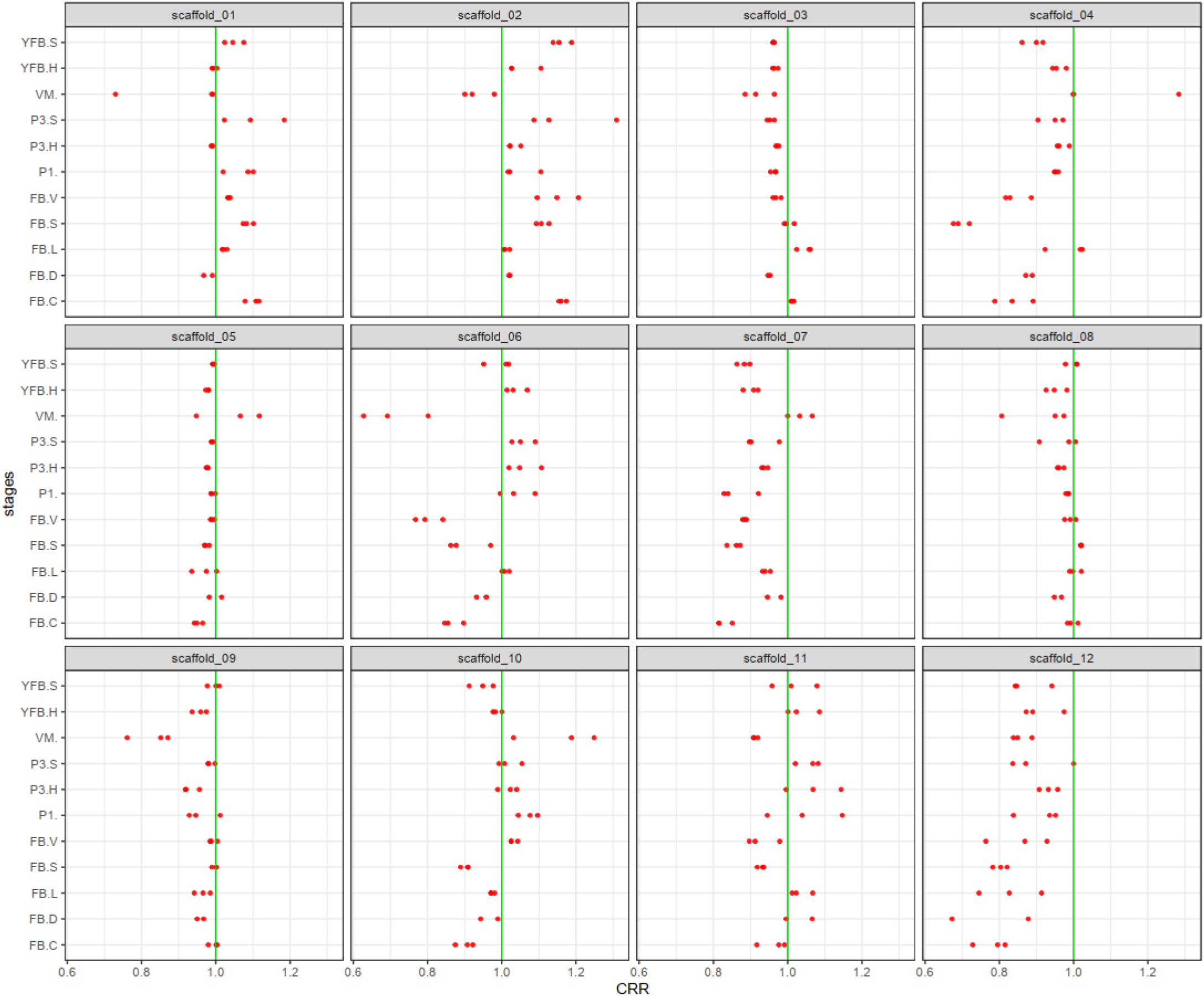
Chromosomal Read Ratio (CRR, the contribution of nuclei to the expression of scaffolds) across stages and tissues. Red dots represent the biological replicates separately. Only scaffold 12 was consistently biased towards PC9, whereas other scaffolds varied depending on developmental stages and tissue types (CRR min-max=0.62-1.31; max difference 37.2%). Values smaller than 1 mean the dominance of the PC9 nucleus over PC15 while the opposite marks the dominance of PC15. Abbreviations as follows: ‘VM’ vegetative mycelium; ‘P1’ stage 1 primordium; ‘P3’ stage3 primordium; ‘YFB’ young fruiting body, ‘FB’ fruiting body; ‘H’ cap (entire); ‘C’ cap trama (only the inner part, without Lamellae, or skin); ‘L’ lamellae; ‘S’ stipe; ‘V’ cuticle; ‘D’ dedifferentiated tissue of cap.

**Figure S11:**
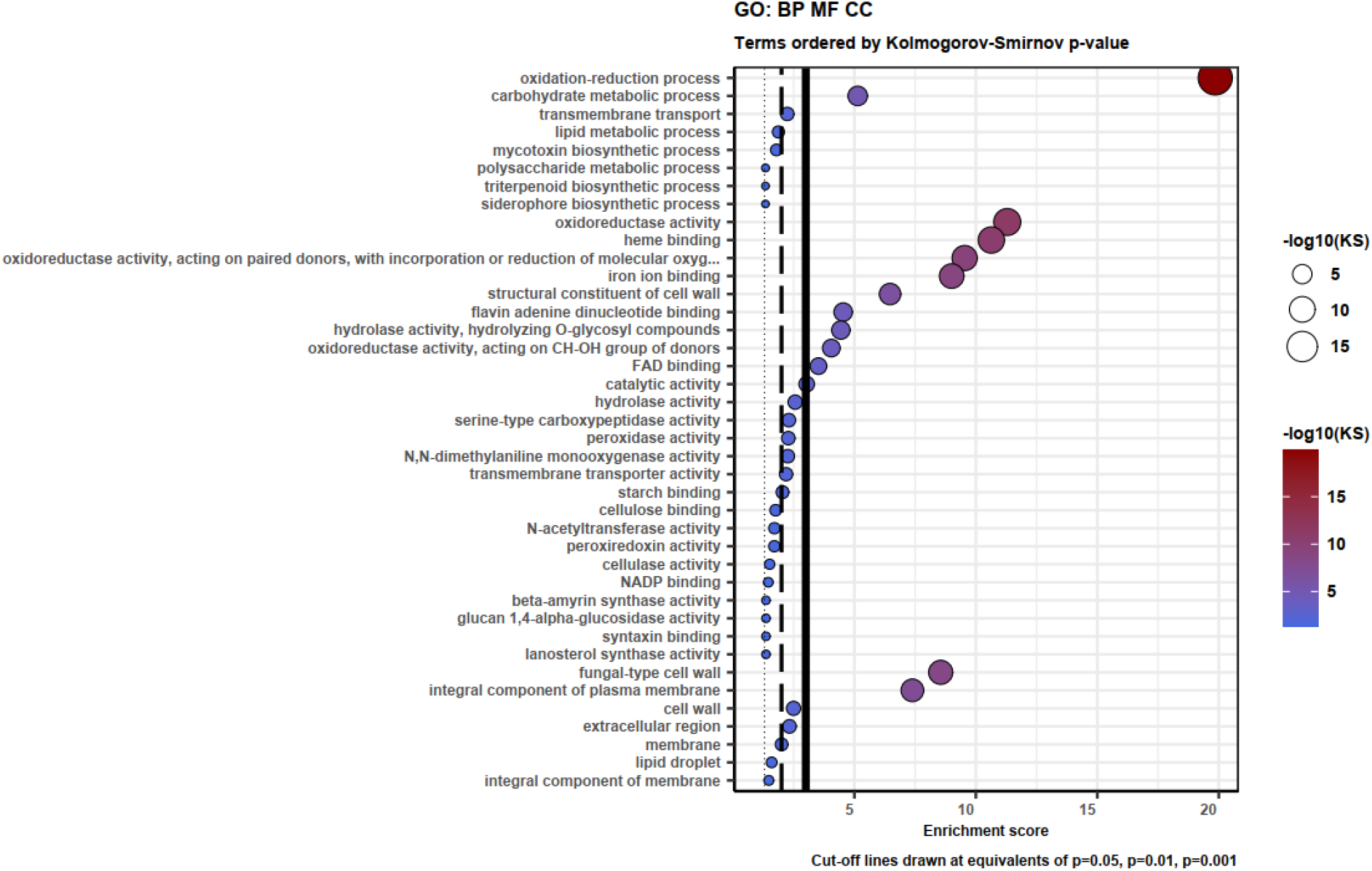
Gene Ontology (GO) enrichment for ASE genes with minimum two-fold change at least in one stage between the two parental nuclei. KS means the p-value of Kolmogorov-Smirnov test implemented in the R package ‘topGO’. BP: Biological Process; MF: Molecular Function; CC: Cellular component.

**Figure S12:**
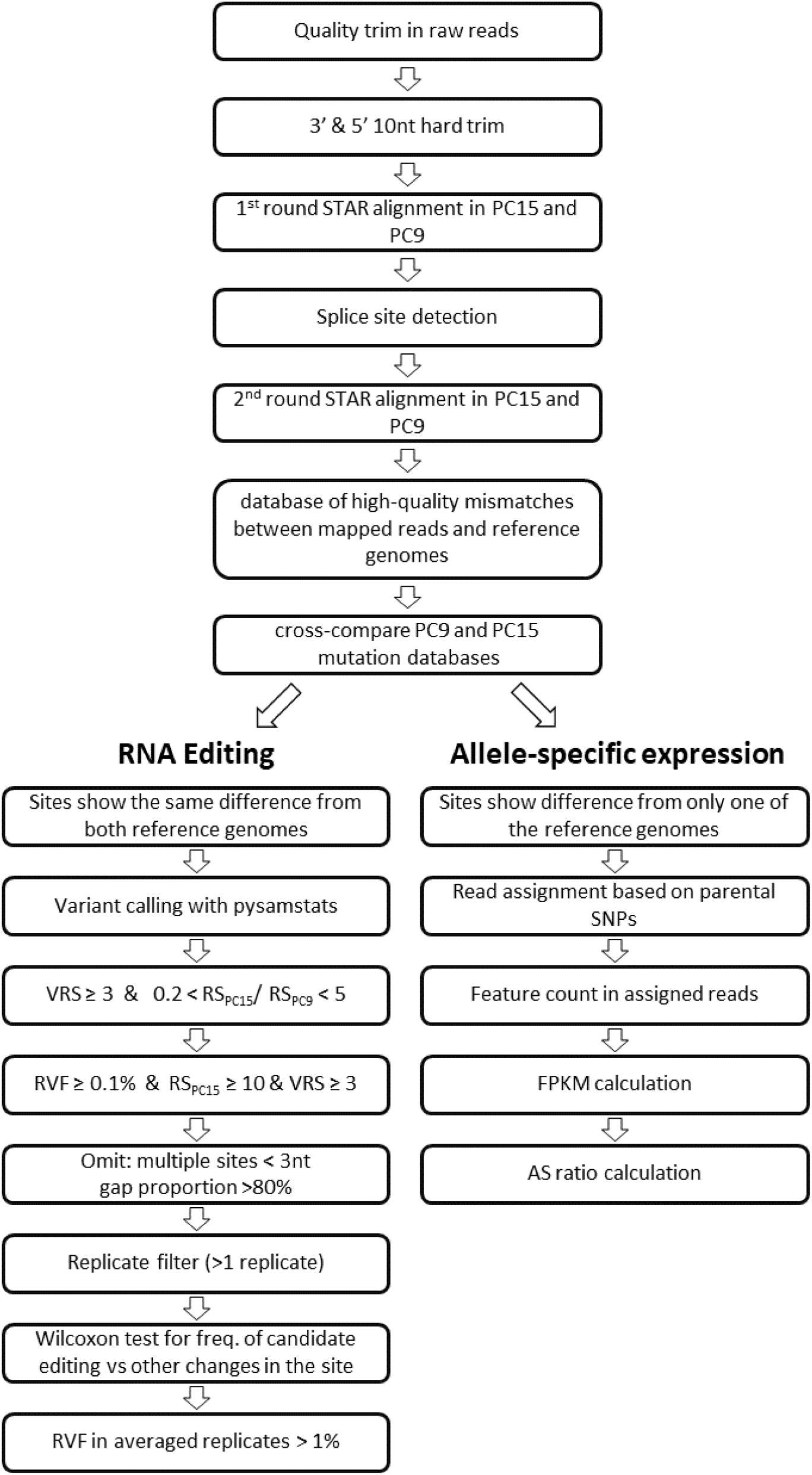
Pipeline of RNA editing and Allele Specific Expression annotation. Abbreviations: VRS Variant Read Support; RS Read Support; RVF Relative Variant Frequency.

**Figure S13:**
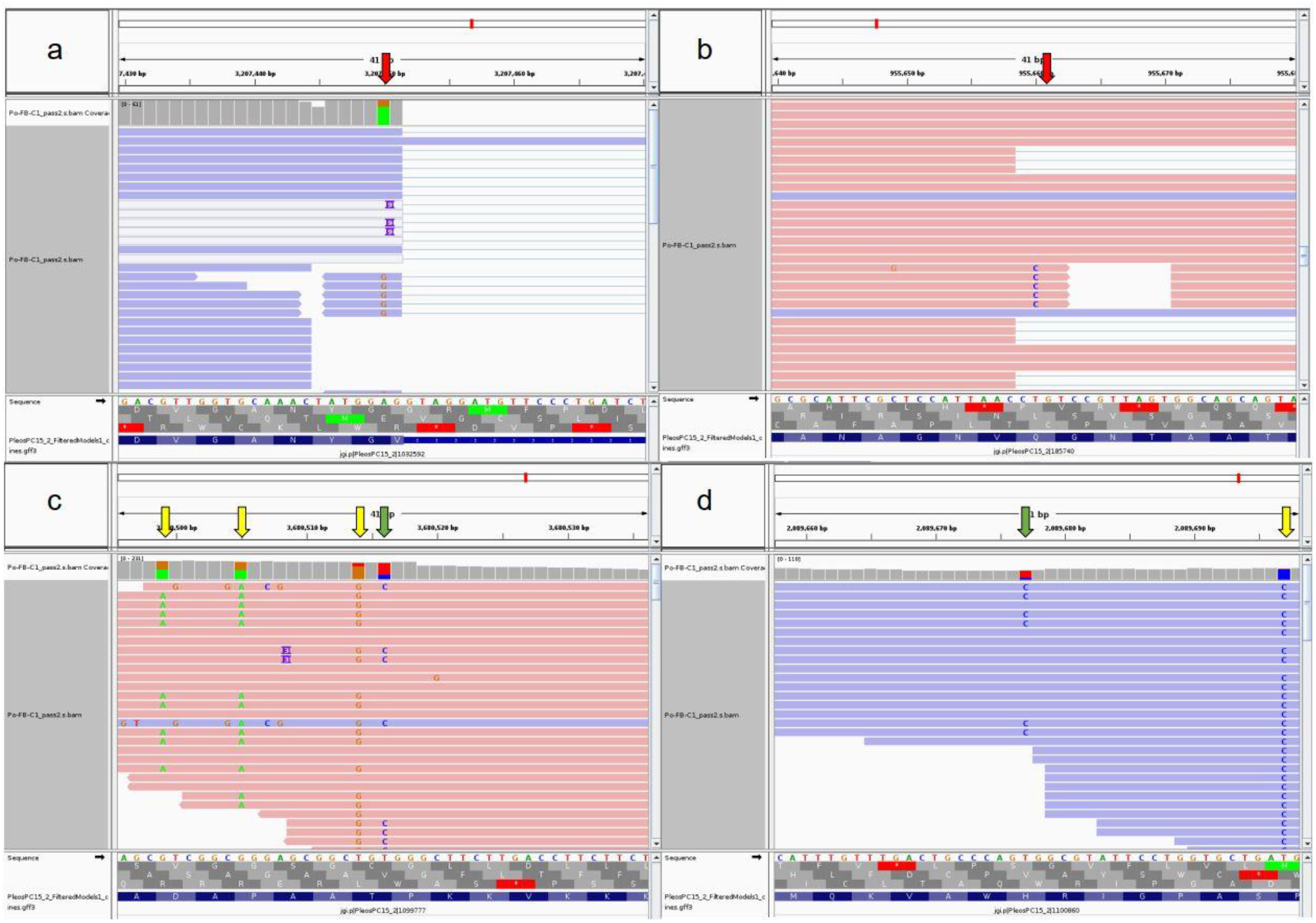
Examples for variants of different types. In a) and b) erroneous read alignment around splice sites causing variants (red arrows) similar to RNA editing. In c) and d) green arrows represent potential RNA editing sites, while yellow arrows represent allele specific SNPs.

**Figure S14:**
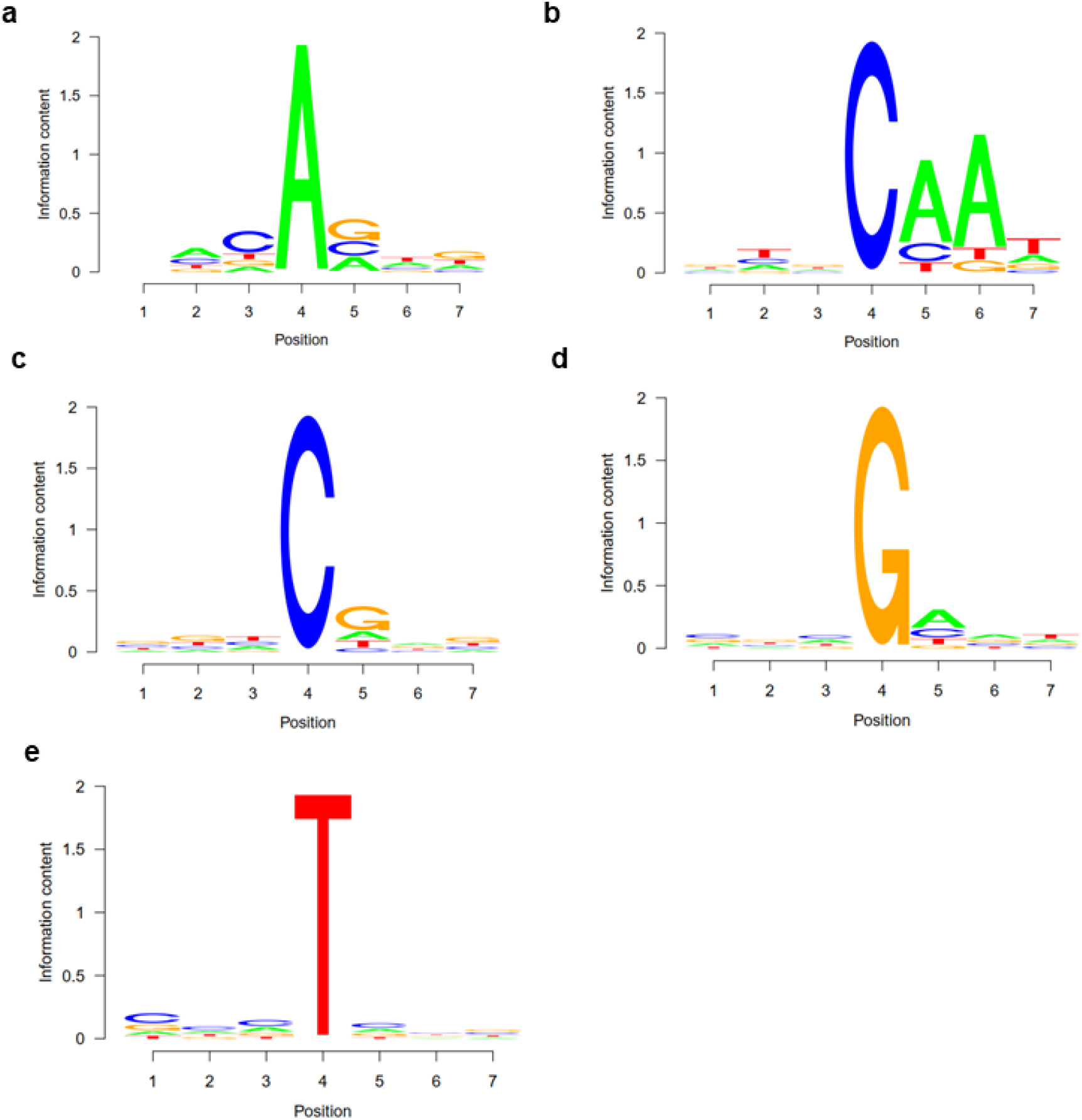
Sequence motifs surrounding the most frequent candidate RNA editing changes displayed as sequence logos. 4^th^ position represents the variants among reads. 1-3 is the upstream 3 positions, while 4-7 is the downstream 3 positions. a) A-to-G b) C-to-A c) C-to-T d) G-to-A e) T-to-C changes.

**Figure S15:**
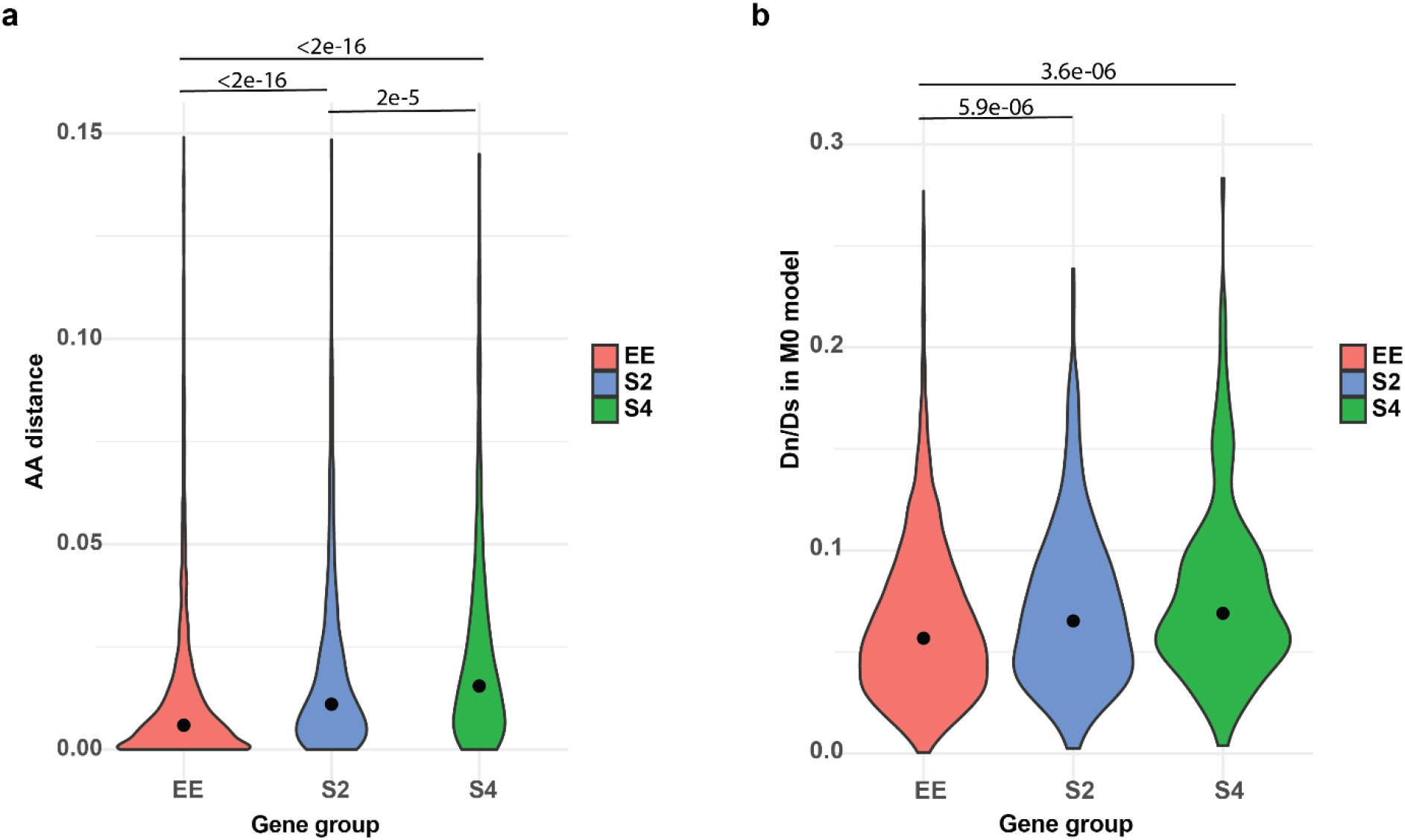
a) Maximum likelihood amino acid distances between PC15 and PC9 proteins (model=WAG). b) dN/dS distribution for the three gene groups under the M0 model in CodeML. Abbreviations: equally expressed genes EE, genes with allele specific expression at least in one stage with two- (S2) or four-fold change (S4). Lines and p-values (<1e-3) represent the pairwise comparison of Nemenyi post-hoc test of Kruskal Wallis test.

**Figure S16:**
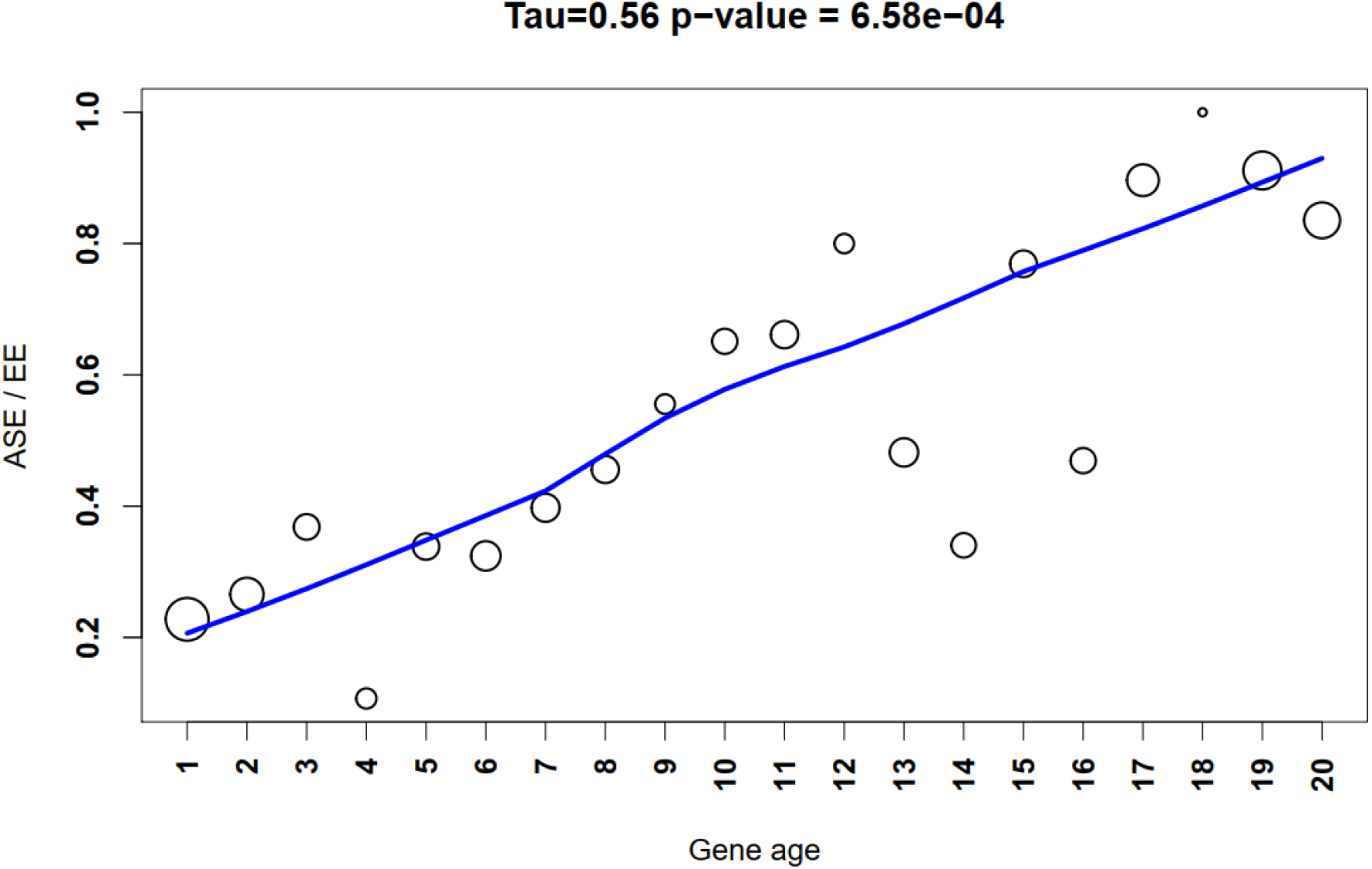
Proportion of Allele Specific Expression (ASE) shows a significant increasing tendency (Mann-Kendall statistics in the title) across gene ages among developmentally regulated genes (Fold Change >=4). Size of circles represents the number of proteins (log10 transformed).

**Figure S17:**
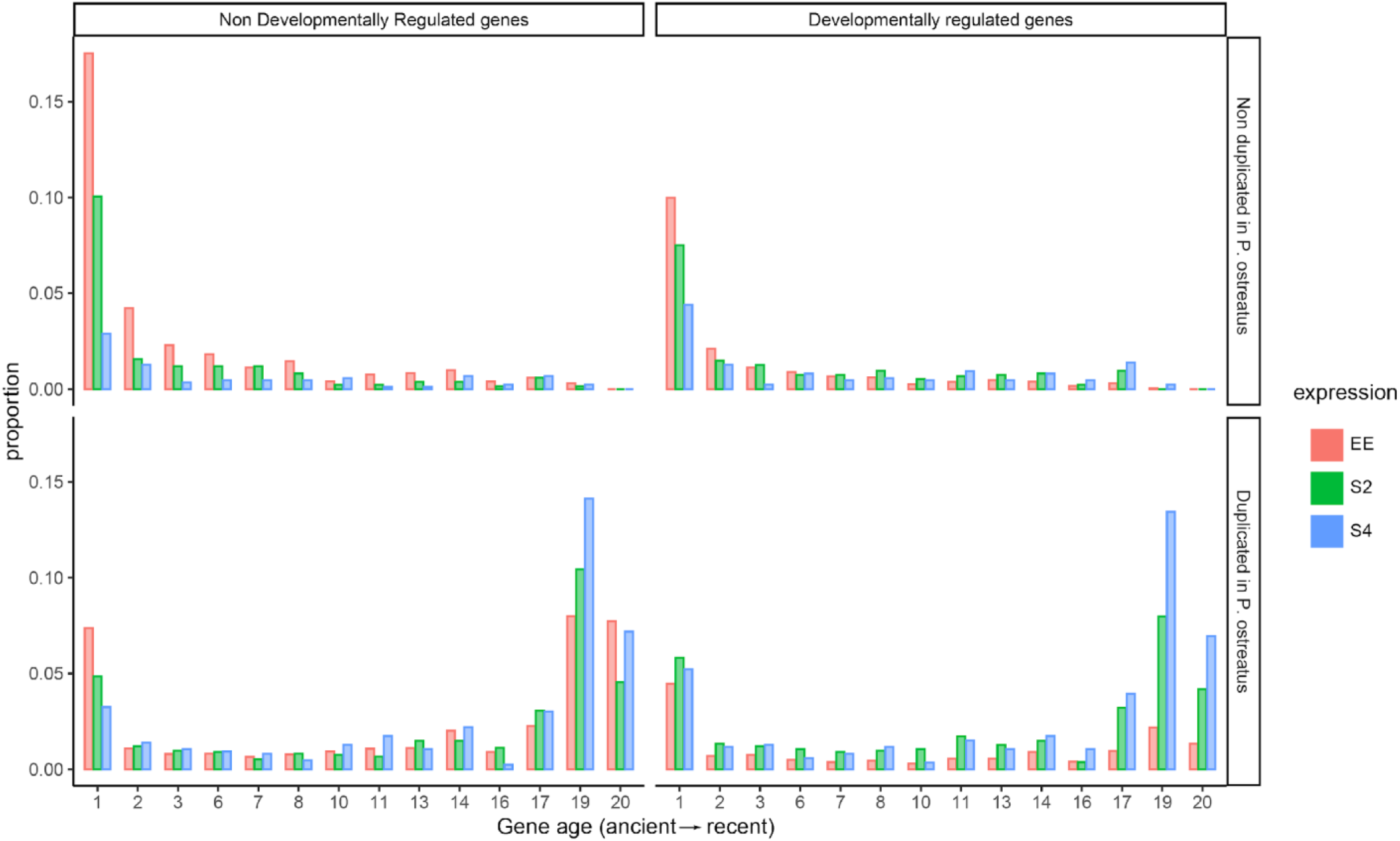
Distribution of genes across gene ages (1 representing oldest and 20 the youngest), broken down by ASE, developmental regulation, and duplication in *Pleurotus ostreatus*. Bars represent proportions of gene numbers relative to the total gene number of a group (equal, S2 or S4). Abbreviations: equal – equally expressed genes; S2 – ASE with two-fold change; S4 –ASE with four-fold change; NDR – non-developmentally regulated gene DR – developmentally regulated gene.

**Figure S18:**
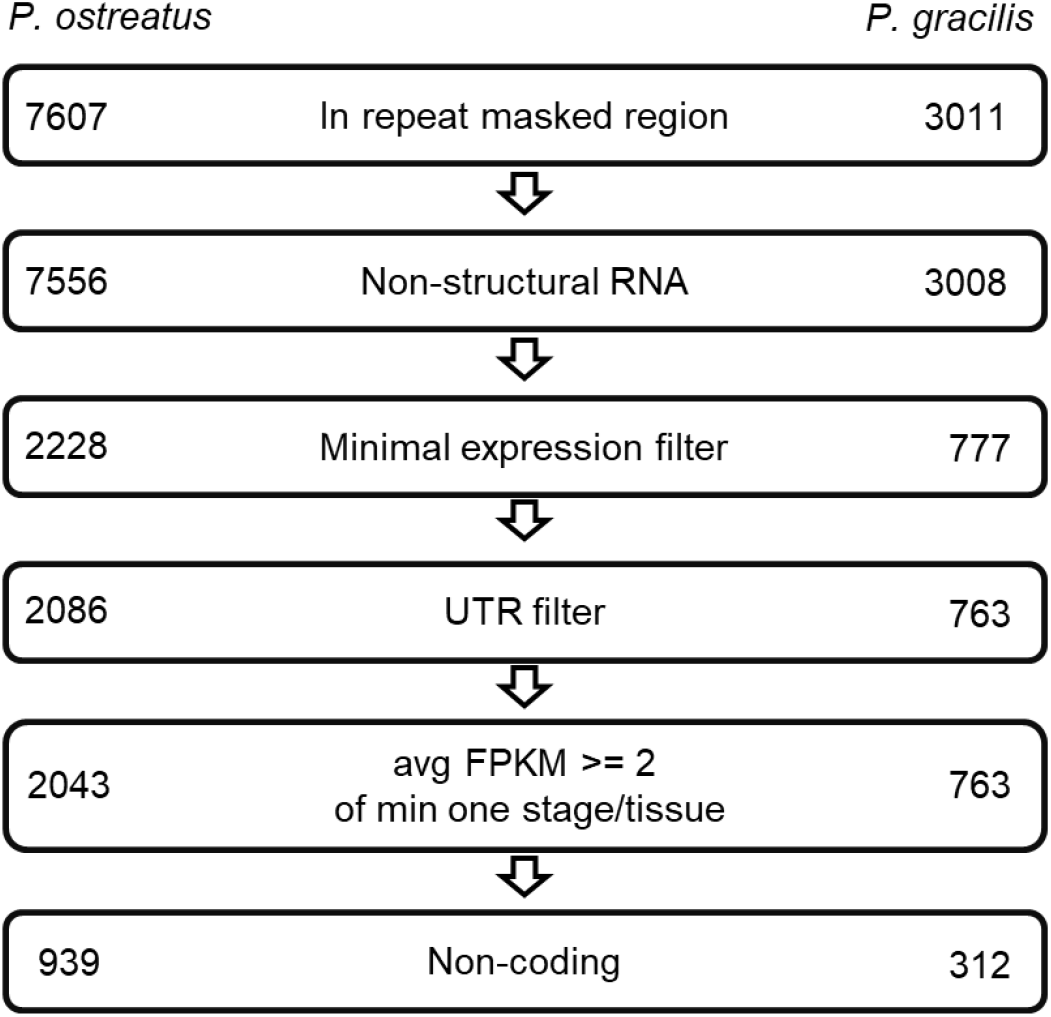
Pipeline of Natural Antisense Transcripts. Numbers represent the retained transcripts in each filtering step.

**Figure S19:**
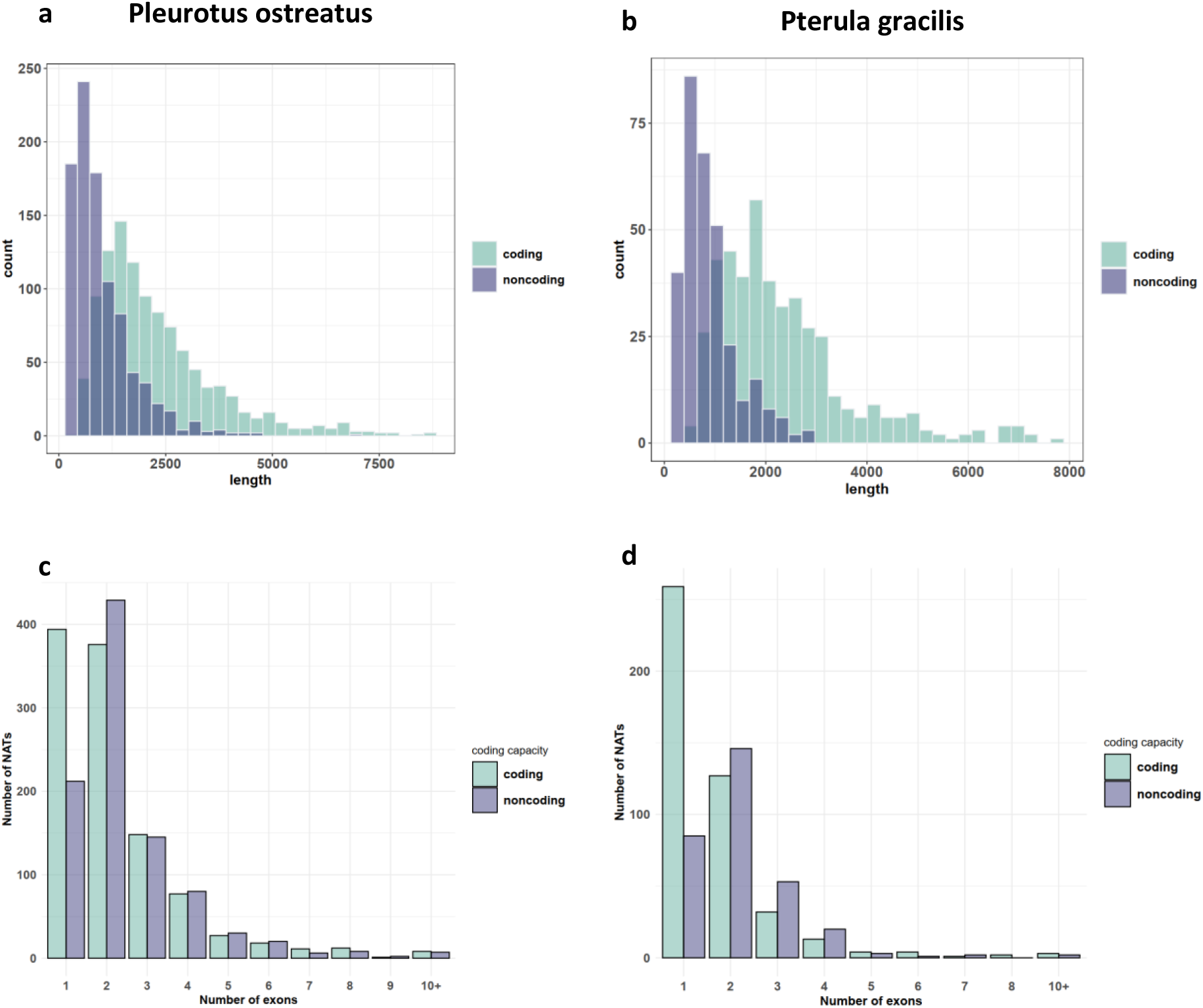
Length (a/b) and exon (c/d) number distribution of natural antisense transcripts in *Pleurotus ostreatus* (a/c) and *Pterula gracilis* (b/d). NATs were divided into putatively coding and noncoding categories using the default settings of CPC2 (Kang et al., 2017)

**Figure S20:**
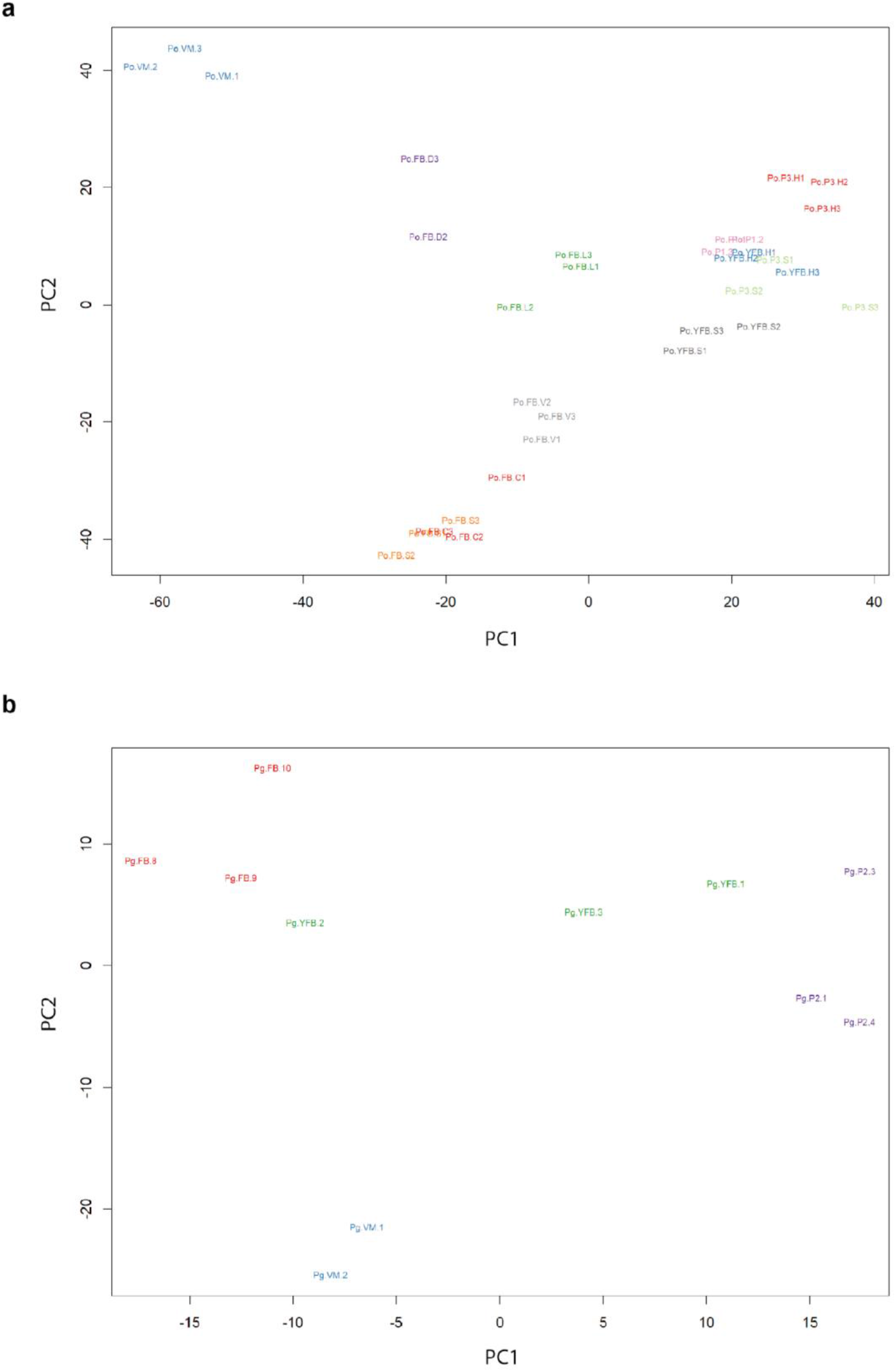
Principal Component Analysis (PCA) plot based on the expression of Natural Antisense Transcripts detected in a) *Pleurotus ostreatus* and b) *Pterula gracilis*. Abbreviations as follows: ‘VM’ vegetative mycelium; ‘P1’ stage 1 primordium; ‘P3’ stage3 primordium; ‘YFB’ young fruiting body, ‘FB’ fruiting body; ‘H’ cap (entire); ‘C’ cap trama (only the inner part, without Lamellae, or skin); ‘L’ lamellae; ‘S’ stipe; ‘V’ cuticle; ‘D’ dedifferentiated tissue of cap.

**Figure S21:**
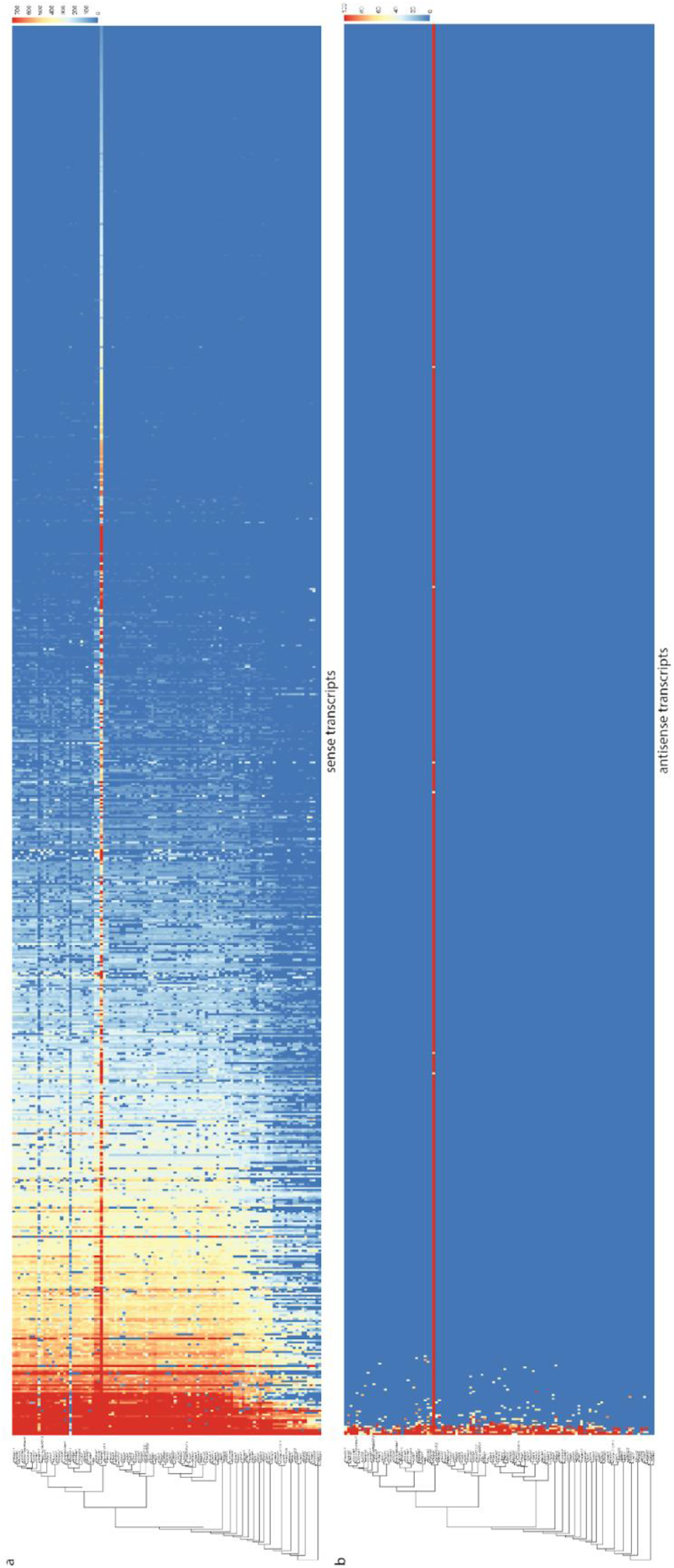
Conservation of sense genes and their antisense transcripts (NAT) in *Pterula gracilis*. a) similarity of sense transcripts measured with –log10(e-value) from MMSeq2 search b) mapping score of antisense transcripts based on Minimap2. Warmer color represents a higher similarity according to the scales

**Figure S22:**
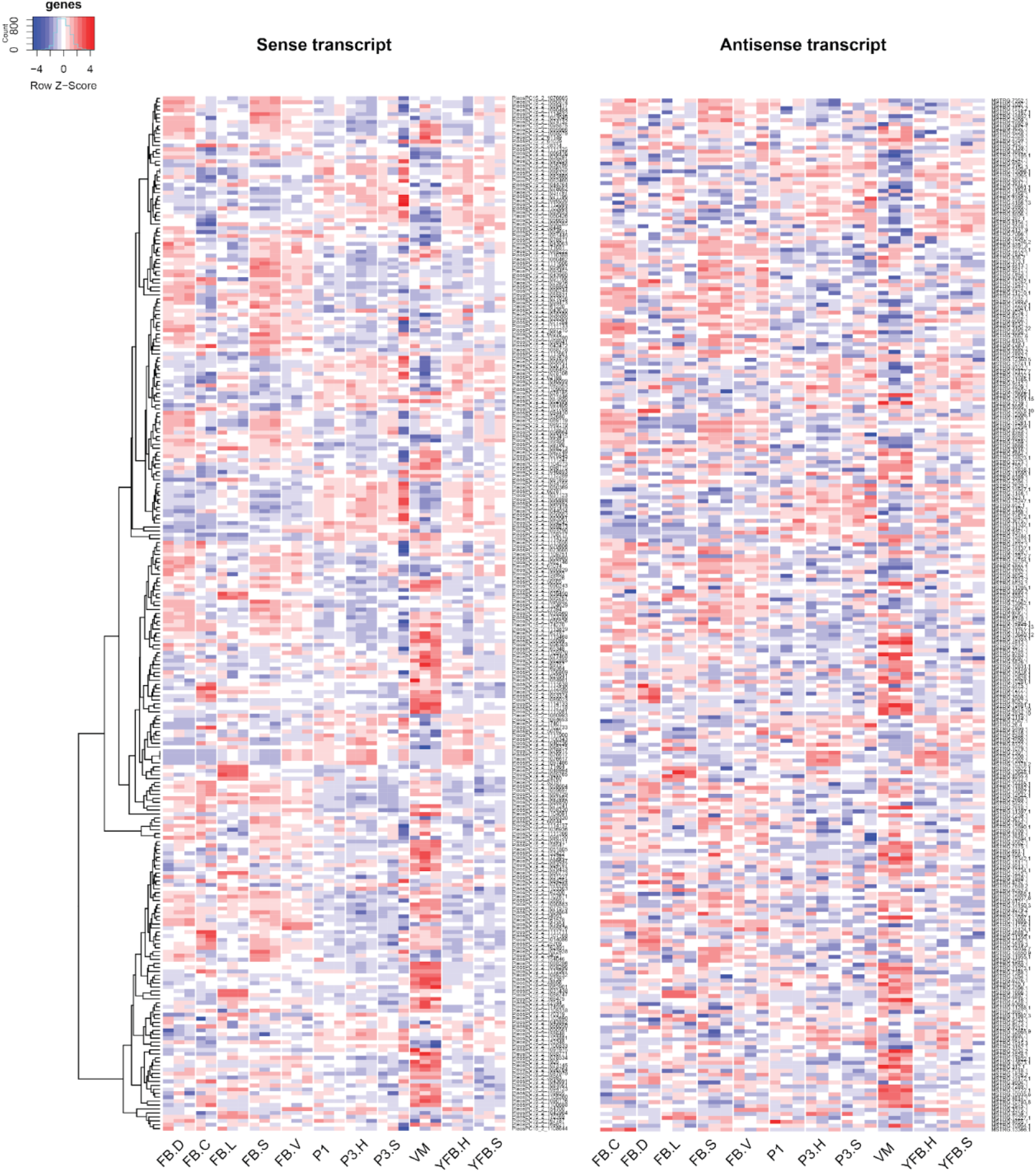
Expression pattern of 263 sense transcripts and their antisense transcripts which showed significant positive correlation (Pearson r ≥ 0.7, p < 0.05) in *Pleurotus ostreatus*. Abbreviations as follows: ‘VM’ vegetative mycelium; ‘P1’ stage 1 primordium; ‘P3’ stage3 primordium; ‘YFB’ young fruiting body, ‘FB’ fruiting body; ‘H’ cap (entire); ‘C’ cap trama (only the inner part, without Lamellae, or skin); ‘L’ lamellae; ‘S’ stipe; ‘V’ cuticle; ‘D’ dedifferentiated tissue of cap.

**Figure S23:**
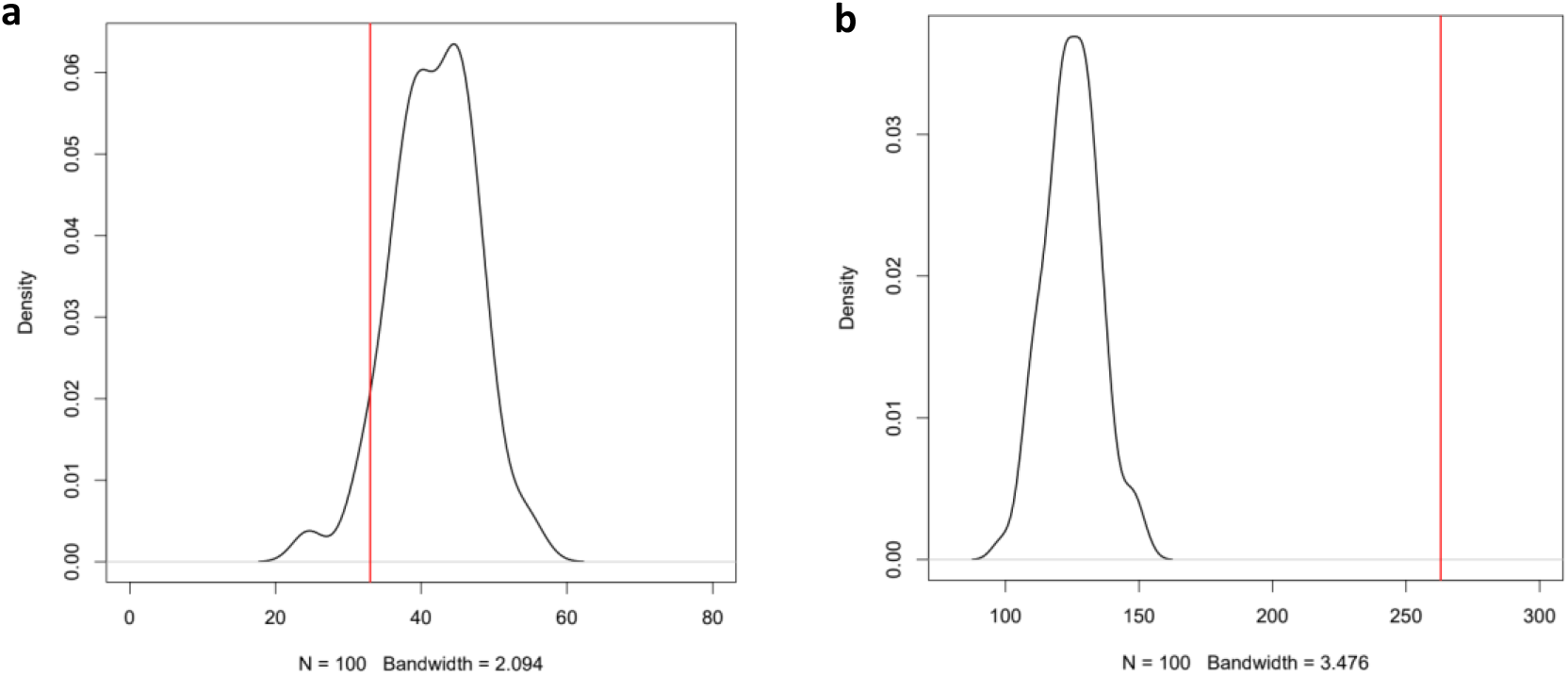
Permutation test for the number of a) negative (r<−0.7, P<0.05) and b) positive (r>0.7, P<0.05) correlations among the expression of NATs and random genes. Red line represents the observed number of significant correlations.

**Figure S24:**
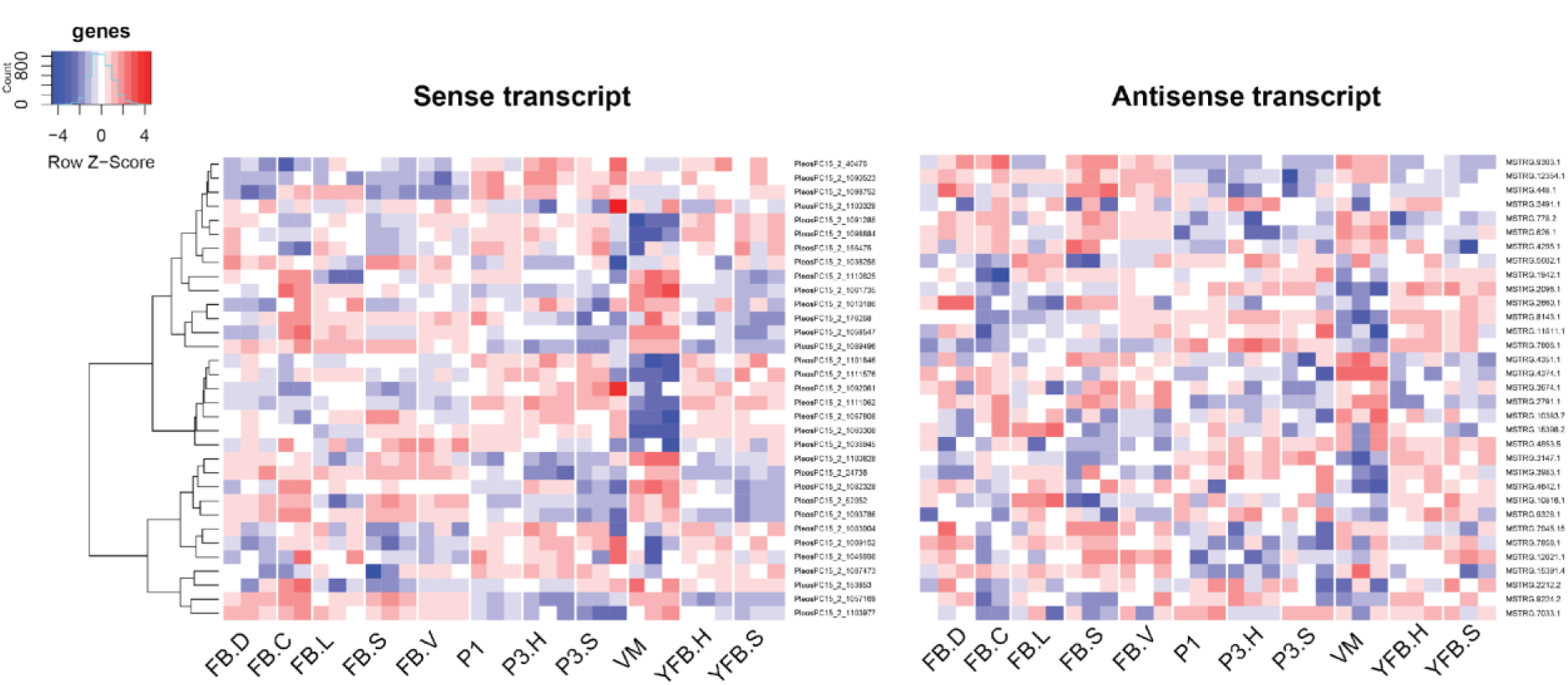
Expression pattern of 33 sense transcripts and their antisense transcripts which showed significant negative correlation (Pearson r ≤ −0.7 P<0.05) in *Pleurotus ostreatus*. Abbreviations as follows: ‘VM’ vegetative mycelium; ‘P1’ stage 1 primordium; ‘P3’ stage3 primordium; ‘YFB’ young fruiting body, ‘FB’ fruiting body; ‘H’ cap (entire); ‘C’ cap trama (only the inner part, without Lamellae, or skin); ‘L’ lamellae; ‘S’ stipe; ‘V’ cuticle; ‘D’ dedifferentiated tissue of cap.

**Figure S25:**
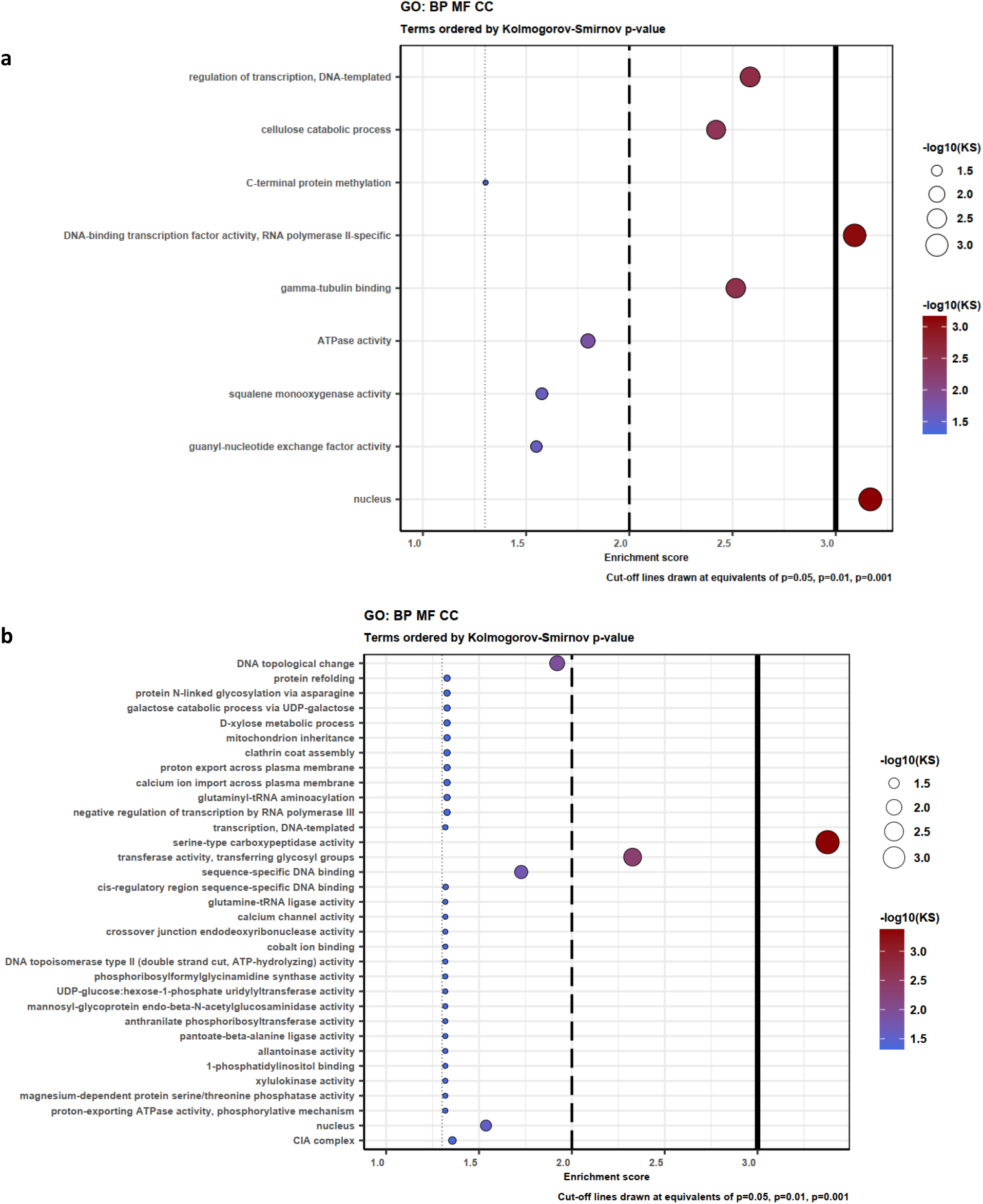
Gene Ontology (GO) enrichment for genes which have Natural Antisense Transcripts a) in *Pleurotus ostreatus* and b) in *Pterula gracilis.* KS means the p-value of Kolmogorov-Smirnov test implemented in the R package ‘topGO’. BP: Biological Process; MF: Molecular Function; CC: Cellular component.

**Figure S26:**
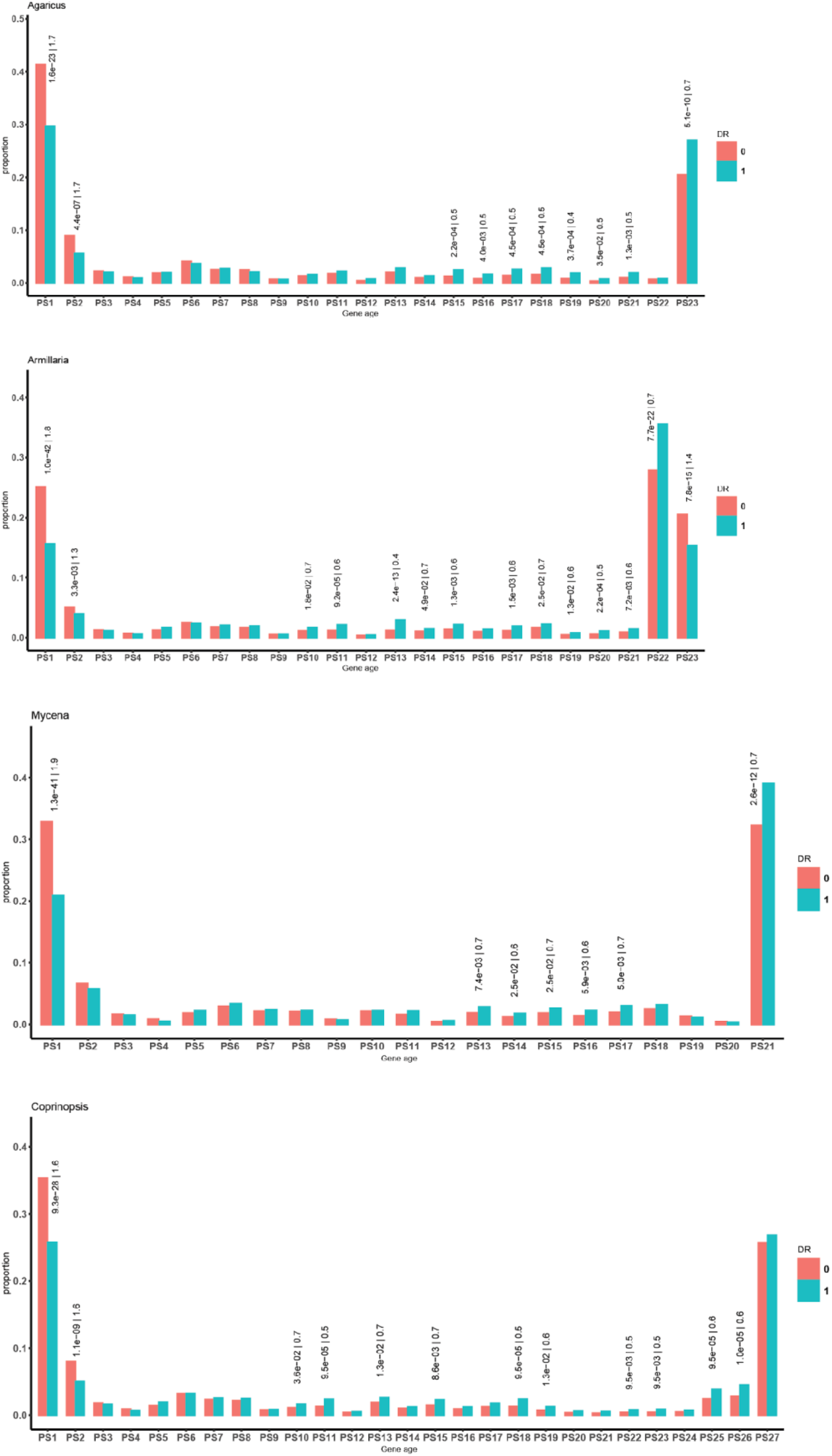
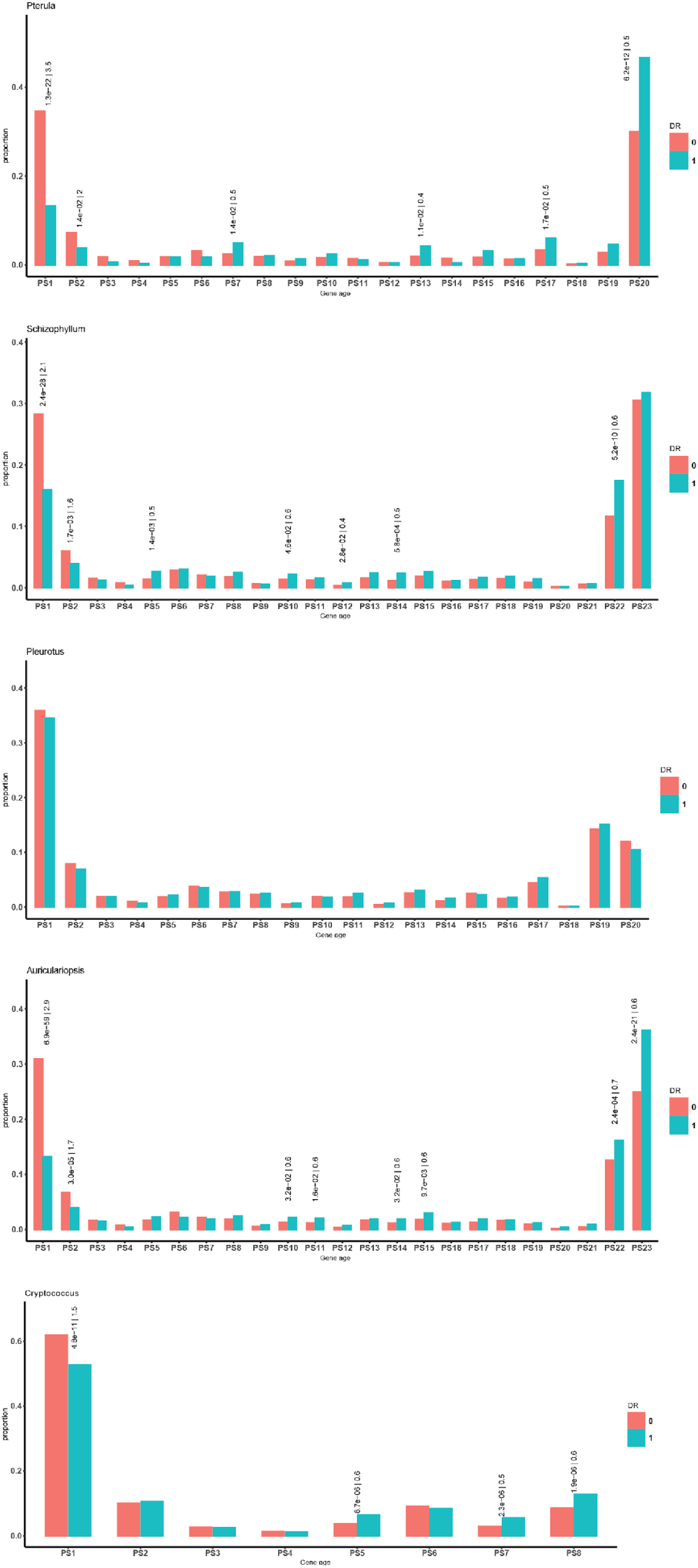
Proportion of Developmentally regulated (DR > 4 FC) genes in different gene ages. P-values and odds ratio above bars comes from Fisher exact test, added only when significant (P<0.05).

**Figure S27:**
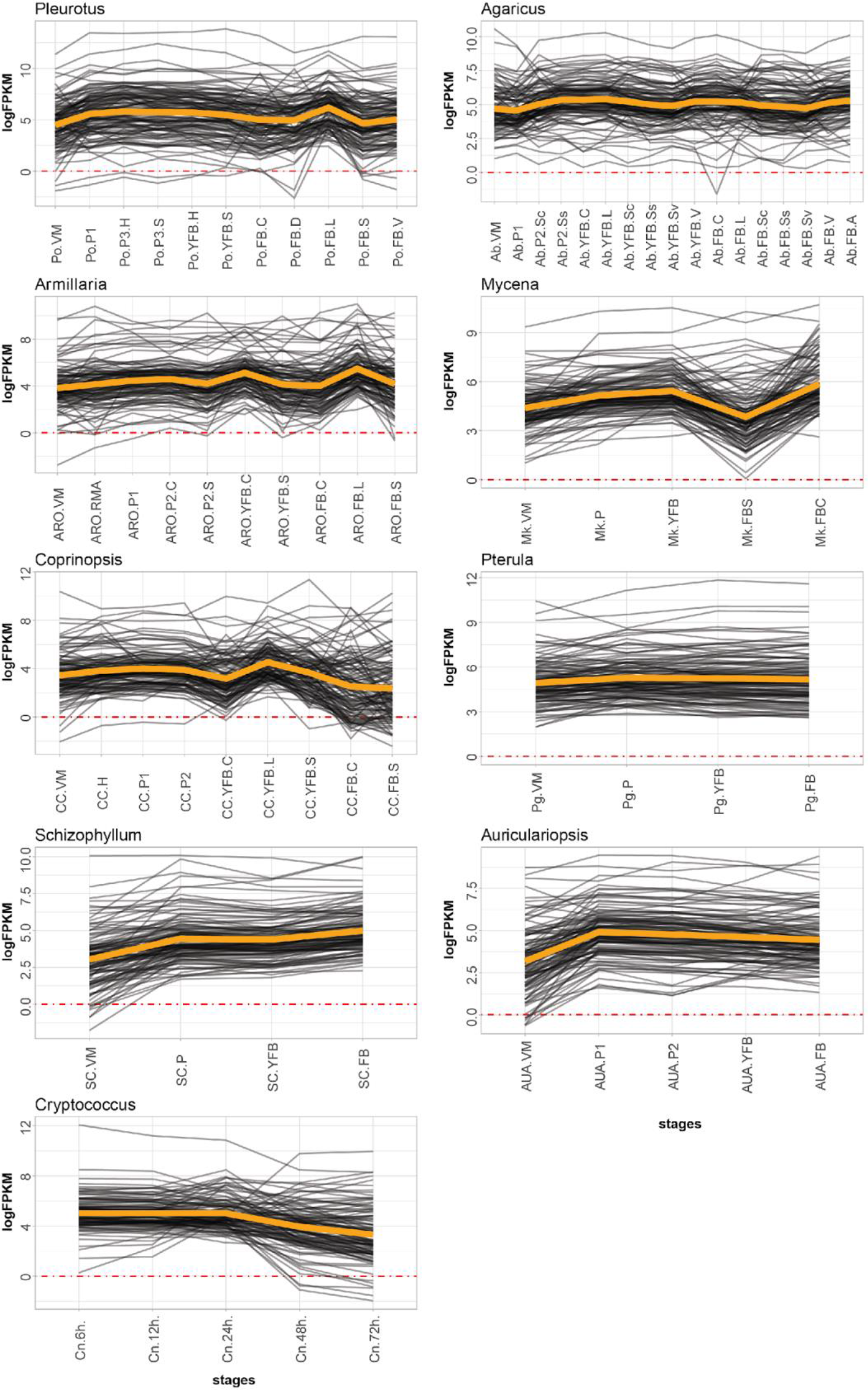
Expression of meiotic genes in the nine species. Abbreviations as follows: ‘VM’ vegetative mycelium; ‘P1’ stage 1 primordium; ‘P3’ stage3 primordium; ‘YFB’ young fruiting body, ‘FB’ fruiting body; ‘H’ cap (entire); ‘C’ cap trama (only the inner part, without Lamellae, or skin); ‘L’ lamellae; ‘S’ stipe; ‘V’ cuticle; ‘D’ dedifferentiated tissue of cap ‘A’ annulus. For stage details see Table S10.

**Figure S28:**
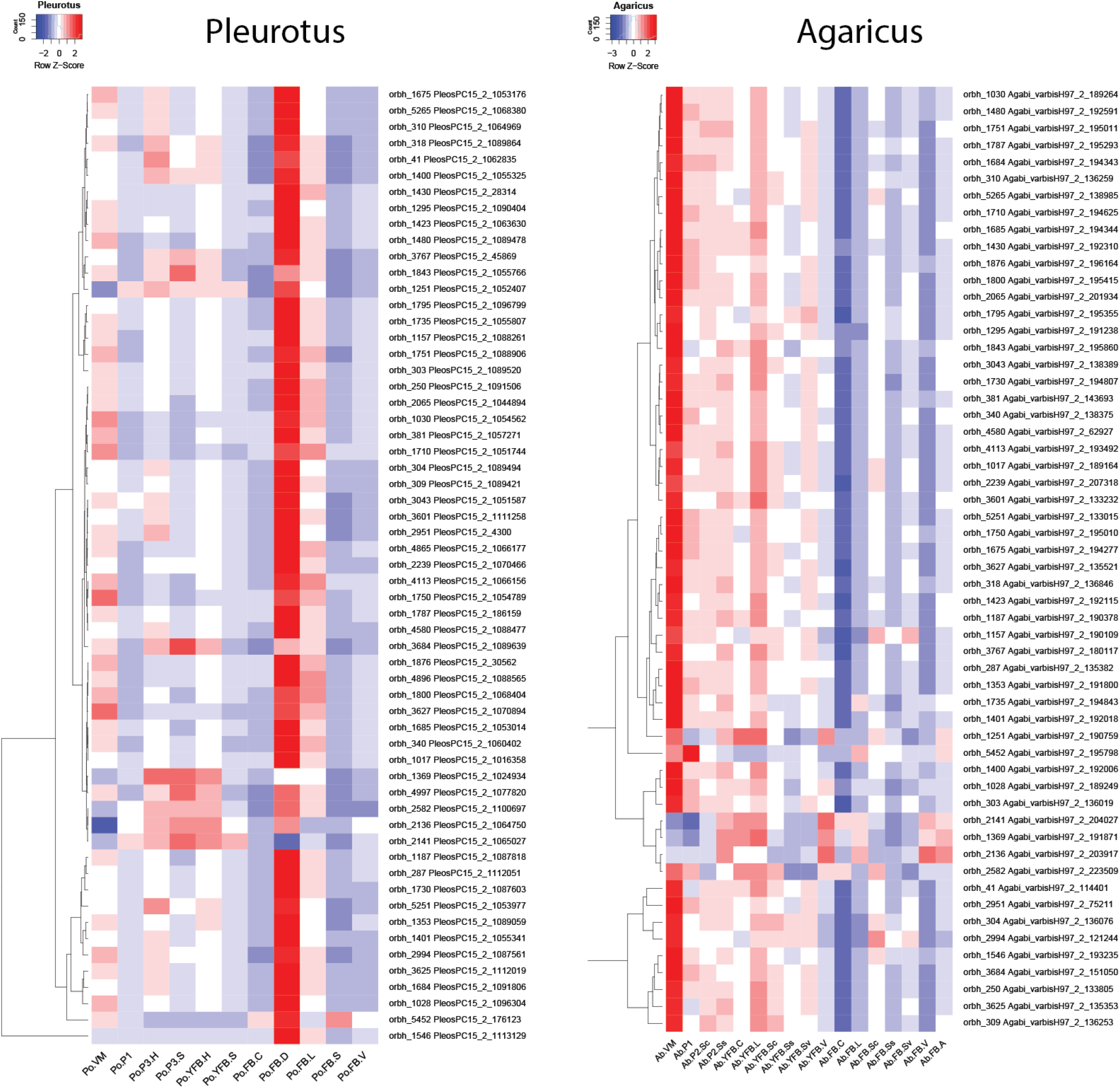
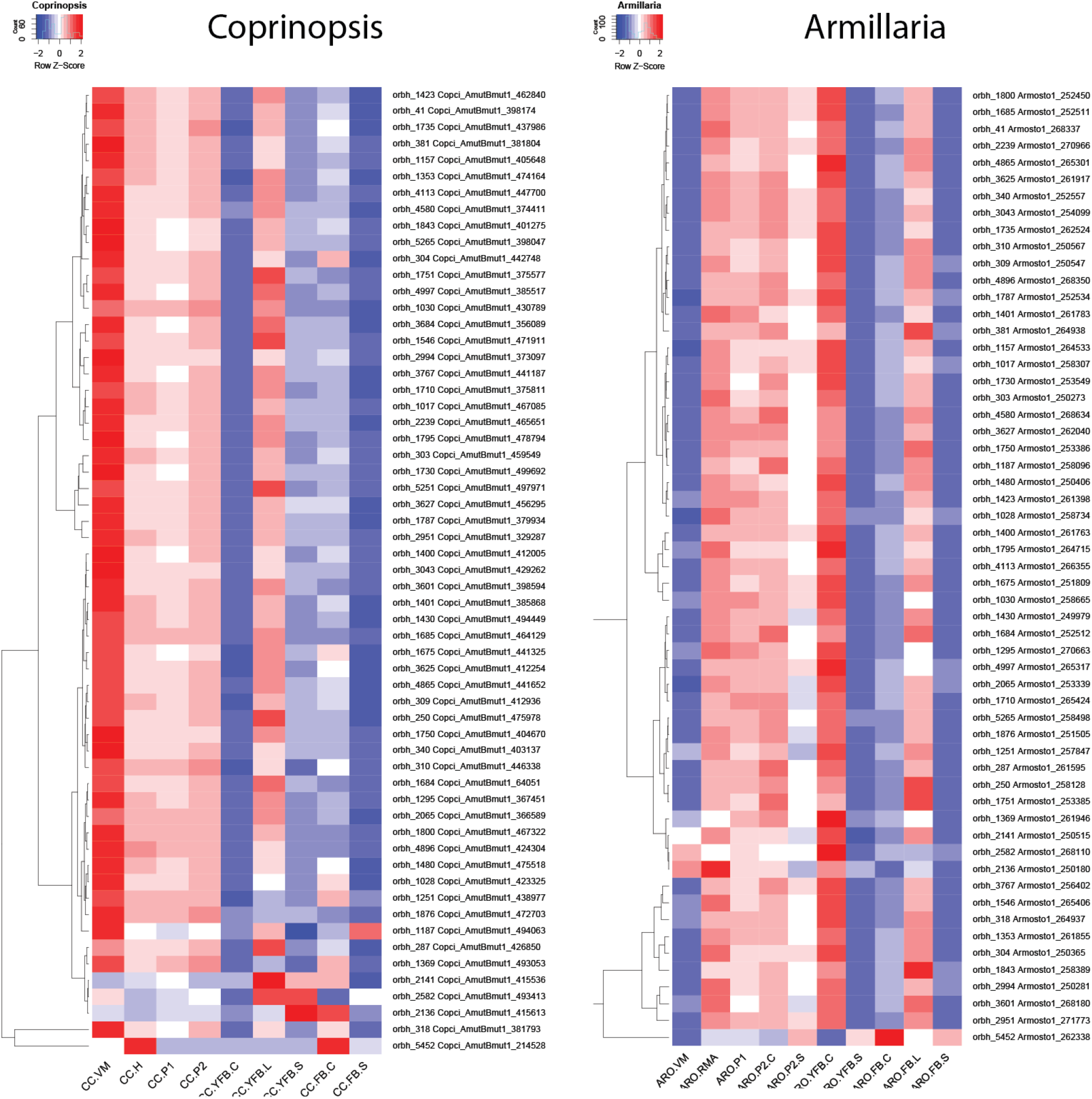
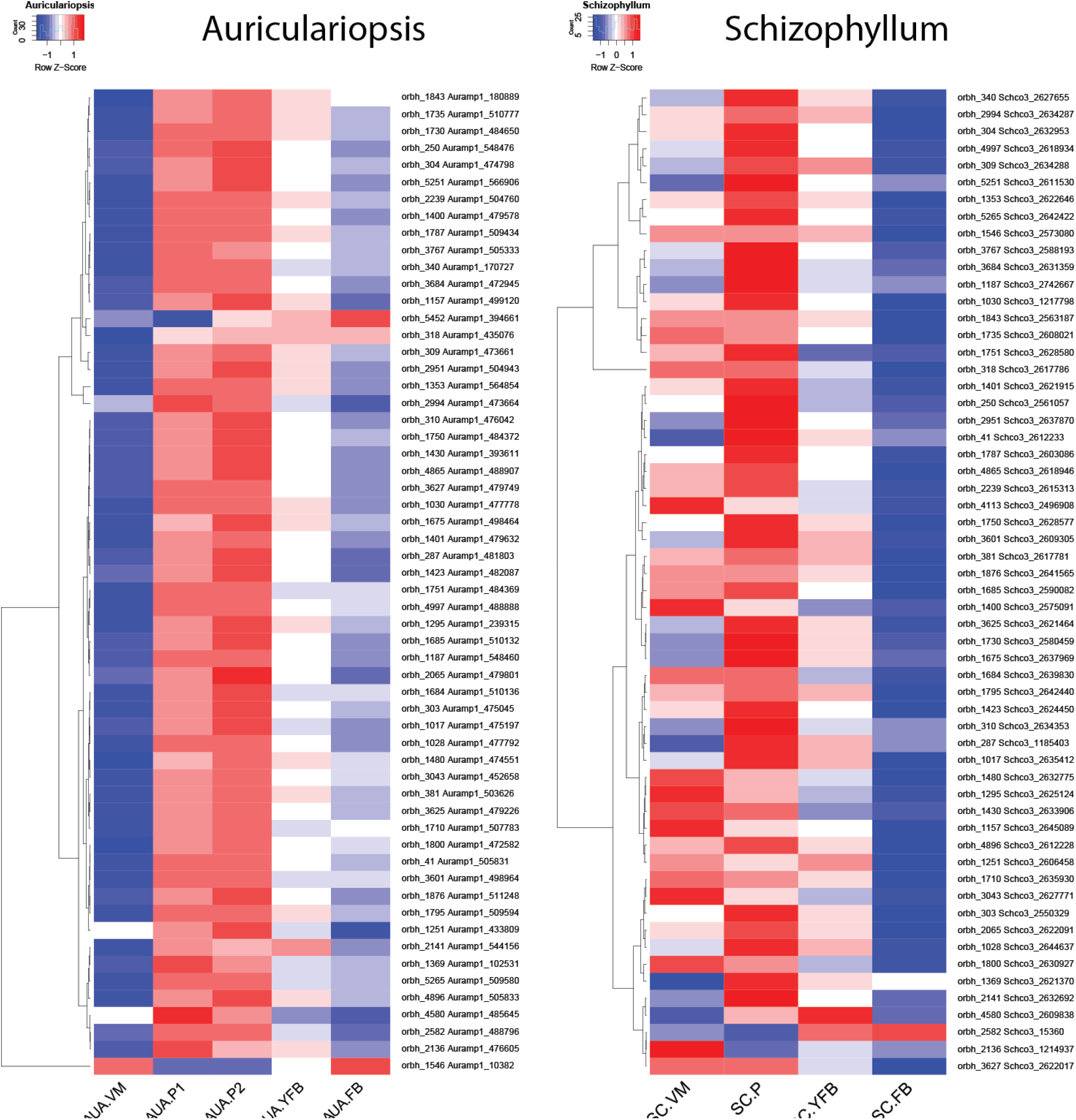
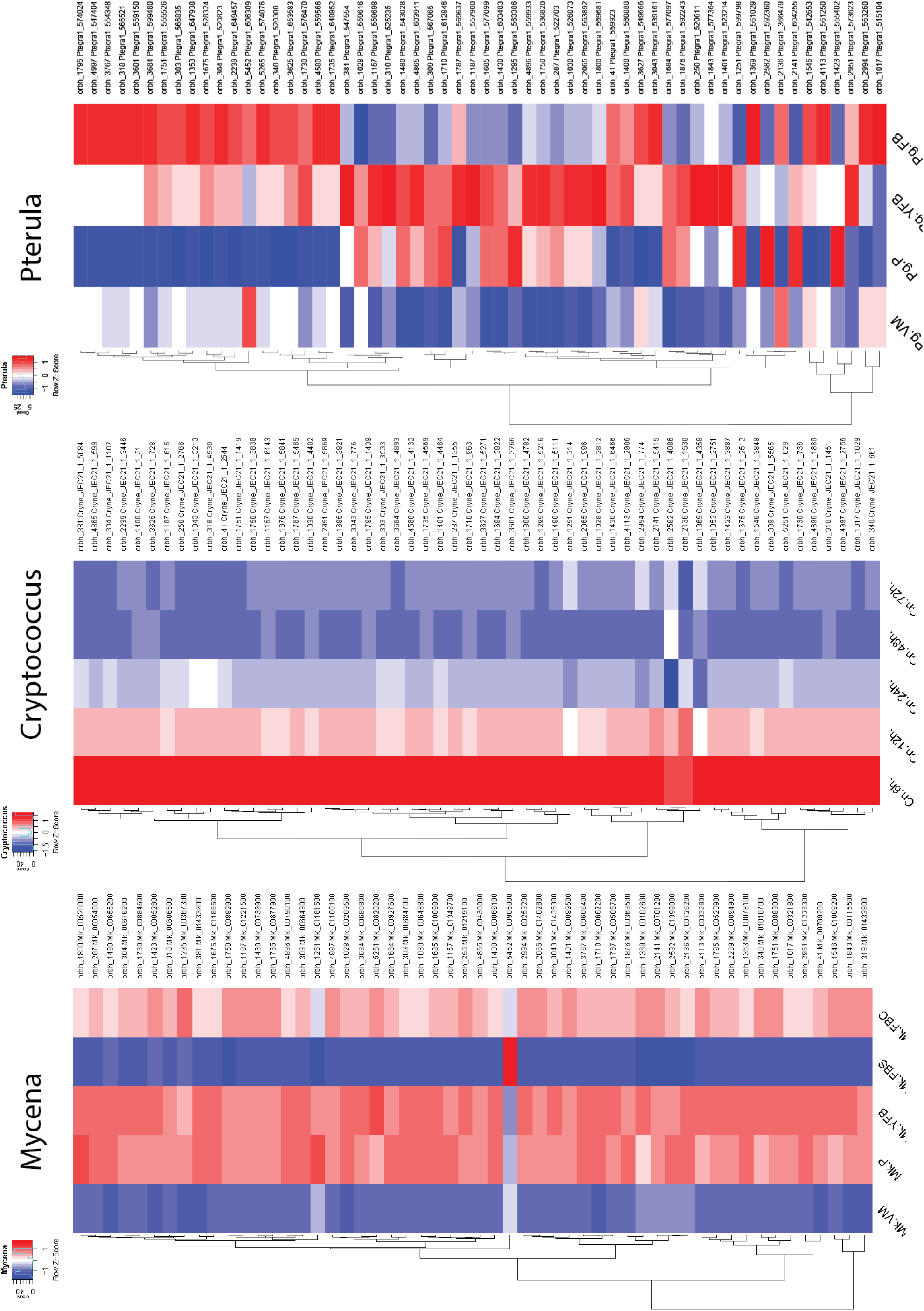
Expression of ribosomal proteins in the nine species. Abbreviations as follows: ‘VM’ vegetative mycelium; ‘P1’ stage 1 primordium; ‘P3’ stage3 primordium; ‘YFB’ young fruiting body, ‘FB’ fruiting body; ‘H’ cap (entire); ‘C’ cap trama (only the inner part, without Lamellae, or skin); ‘L’ lamellae; ‘S’ stipe; ‘V’ cuticle; ‘D’ dedifferentiated tissue of cap ‘A’ annulus. For stage details see Table S10.

**Figure S29:**
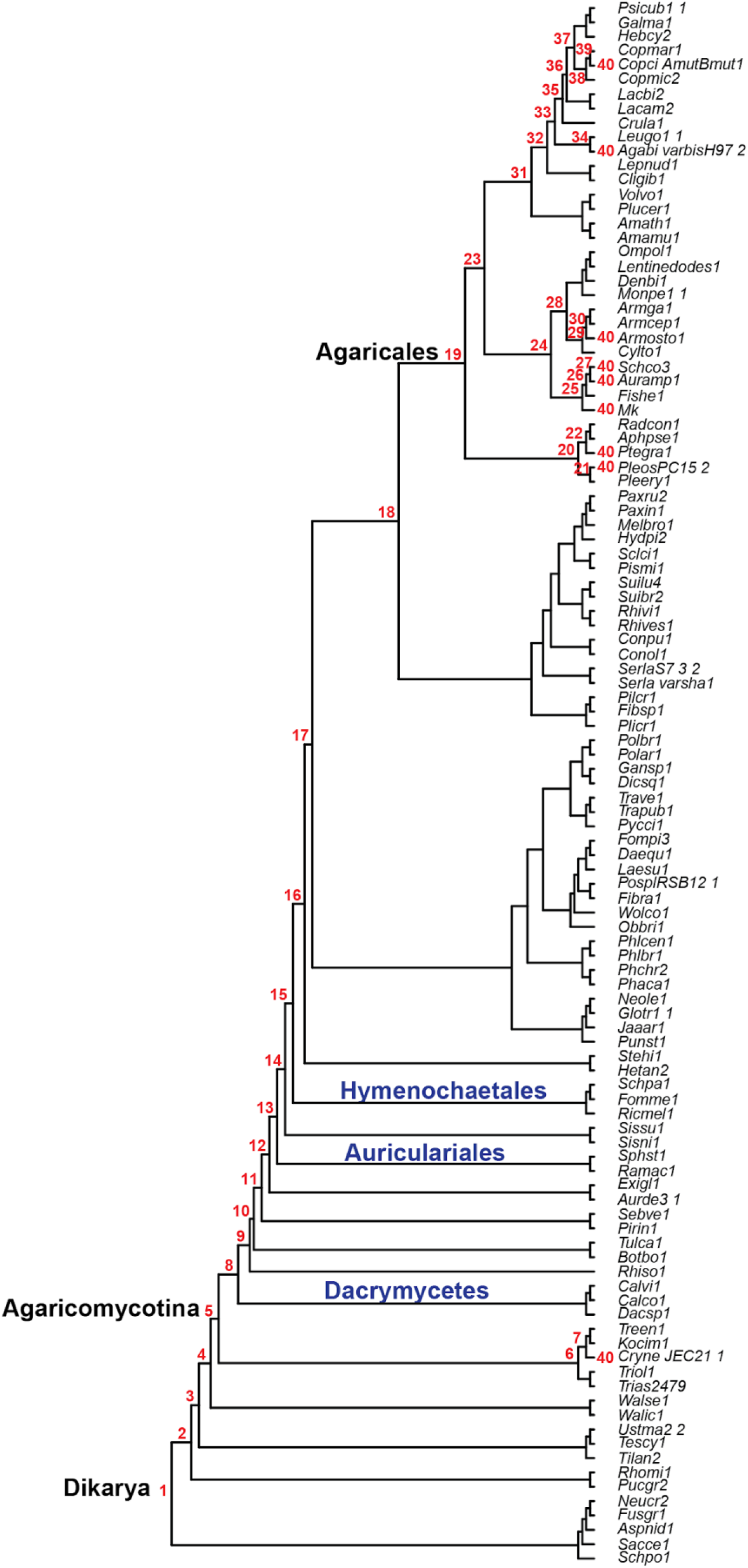
Species tree with the number of relative gene ages (red). For deciphering species IDs see Table S11.

## Captions for Table S1 to S12

**Table S1. (separate file).** Mapping statistics for *Pleurotus ostreatus* and *Pterula gracilis*

**Table S2. (separate file).** Metadata of the 138 FBDR hotspots

**Table S3. (separate file).** Mapped reads to the parental reference genomes and distribution of reads according to parental assignment

**Table S4. (separate file).** Averaged AS ratio among replicates of 4943 genes, which showed imbalance at least one sample.

**Table S5. (separate file).** InterPro domain enrichment analysis for genes with Allele Specific Expression (S2 and S4) among all IPR annotated genes.

**Table S6 (separate file).** Gene Ontology enrichment for genes with Allele Specific Expression.

**Table S7 (separate file).** Detected potential RNA editing sites

**Table S8 (separate file).** Conserved 1:1 ortholog groups.

**Table S9 (separate file).** Putative defense related ortholog groups.

**Table S10 (separate file).** Species which were used for transcriptomic analysis

**Table S11 (separate file).** Species, which were used for the species tree estimation and phylostratigraphy

**Table S12 (separate file).** Gene Ontology enrichment for CM-specific and Shared orthologs

